# Perivascular adipose tissue remodeling impairs vasoreactivity in thermoneutral-housed rats

**DOI:** 10.1101/2024.05.09.593330

**Authors:** Melissa M Henckel, Ji Hye Chun, Leslie A Knaub, Gregory B Pott, Georgia E. James, Kendall S Hunter, Robin Shandas, Lori A Walker, Jane E-B Reusch, Amy C Keller

## Abstract

**Objective:** Vascular pathology, characterized by impaired vasoreactivity and mitochondrial respiration, differs between the sexes. Housing rats under thermoneutral (TN) conditions causes vascular dysfunction and perturbed metabolism. We hypothesized that perivascular adipose tissue (PVAT), a vasoregulatory adipose depot with brown adipose tissue (BAT) phenotype, remodels to a white adipose (WAT) phenotype in rats housed at TN, driving diminished vasoreactivity in a sex-dependent manner.

**Methods:** Male and female Wistar rats were housed at either room temperature (RT) or TN. Endpoints included changes in PVAT morphology, vasoreactivity in vessels with intact PVAT or transferred to PVAT of the oppositely-housed animal, vessel stiffness, vessel mitochondrial respiration and cellular signaling.

**Results:** Remodeling of PVAT was observed in rats housed at TN; animals in this environment showed PVAT whitening and displayed diminished aortae vasodilation (p<0.05), different between the sexes. Juxtaposing PVAT from RT rats onto aortae from TN rats in females corrected vasodilation (p<0.05); this did not occur in males. In aortae of all animals housed at TN, mitochondrial respiration was significantly diminished in lipid substrate experiments (p<0.05), and there was significantly less expression of peNOS (p<0.001).

**Conclusions:** These data are consistent with TN-induced remodeling of PVAT, notably associated with sex-specific blunting of vasoreactivity, diminished mitochondrial respiration, and altered cellular signaling.

## Introduction

Cardiovascular disease (CVD) is the world’s leading cause of morbidity and mortality(1). It is well known that sex as a biological variable influences CVD risk and pathology (2, 3), with diabetes and smoking conferring different risk in women versus men, further influenced by menopausal status (4). There are also sex differences in CVD symptoms and pathology (4, 5), with women presenting with coronary artery disease differently than men (4). In the vasculature, age-related endothelial dysfunction is different between men and women (6). For example, vascular function has been shown to decrease across the menopause transition in women (7), potentially due to the impact of declining sex hormones on nitric oxide production and bioavailability (6). As there are substantial sex differences in risk factors and disease progression in CVD, it is critical to delineate sex differences in vascular pathology.

Adipose tissue has a strong role in regulating metabolism and cardiometabolic health (8, 9). Aortic perivascular adipose tissue (PVAT) has features of a brown adipose tissue (BAT) depot. PVAT surrounds the vasculature and regulates vascular tone in a paracrine manner by secreting adiponectin, activating eNOS, and/or by secreting NO itself via local eNOS activation (10–17). When animal models are exposed to metabolic stress (i.e. overnutrition, hypertension), PVAT modulation of vascular tone is impaired in a sex-dependent manner, worse in males (18, 19). This metabolic stress causes PVAT remodeling to resemble a white adipose tissue (WAT) phenotype and contributes to endothelial dysfunction (14). Dampened endothelial function is strongly associated with abnormal vasoreactivity and CVD (11, 14). In humans, PVAT quantity has been correlated with decreased insulin sensitivity and diminished regulation of vasoreactivity (20, 21). PVAT in men resembles WAT phenotype more than in women (21), and *in vivo* pathological PVAT phenotype and vascular dysfunction also differs between the sexes (22). In PVAT experiments with spontaneously hypertensive rats, male rats showed a greater change in PVAT phenotype and resultant vascular dysfunction than females (22); this was shown to be from elevated PVAT production of the vasoconstrictor resistin in males (22). In another study, flow cytometry of PVAT immune cell populations described different compositions between male and female Sprague-Dawley rats (23), highlighting the role of PVAT cellular composition in immune contributions to vasoreactivity. Despite these studies, very little is known about how PVAT remodeling impacts vascular reactivity and function.

Rodents housed at TN conditions show measurable differences in physiology (24–26). We previously reported that housing rats at TN caused diminished vascular function in a sex-dependent manner, showing this housing environment as a straightforward way to induce endothelial dysfunction (27). We hypothesized that PVAT remodeling from predominant beige to white alters both vascular physiology and mitochondrial substrate utilization in a sex-specific manner.

## Methods and Materials

### Reagents

Western blotting gels were from BioRad and PVDF membranes were from Millipore. Collagenase, ethylenediaminetetraacetic acid (EDTA), ethylene glycol tetraacetic acid (EGTA), sodium pyrophosphate, sodium orthovanadate, sodium fluoride, okadaic acid, 1% protease inhibitor cocktail, dithiothreitol, magnesium chloride, K-lactobionate, taurine, potassium phosphate, HEPES, digitonin, pyruvate, malic acid, glutamic acid, adenosine diphosphate, succinic acid, oligomycin, carbonyl cyanide 4 (trifluoromethoxy)phenylhydrazone (FCCP), antibody to β-actin (mouse), phenylephrine and acetylcholine, trypsin inhibitor and cytochrome c were procured from Sigma-Aldrich (MO, USA). Dimethyl sulfoxide (DMSO), sodium chloride, sucrose, CitriSolv, and bovine serum albumin were purchased from Fisher Scientific (PA, USA). S-Nitroso-N-Acetyl-D,L-Penicillamine (SNAP) was procured from Cayman Chemical (MI, USA).

### Antibodies

Antibodies to total endothelial nitric oxide synthase (eNOS, Cell Signaling #9572S, 1:500, mouse), and Ser1177 phosphorylated eNOS (peNOS, Cell Signaling #9571S, 1:500 Rabbit), were obtained from Cell Signaling (MA, USA). For the ratio of phosphorylated to total protein, alternate host animal antibodies and alternate secondary antibodies were used with different wavelengths, to eliminate the possibility of signal bleed-through. Antibody cocktail to representative subunits of mitochondrial oxidative phosphorylation (Total OXPHOS Rodent WB Antibody Cocktail Abcam #ab110413,1:1500, mouse) complexes I (subunit NDUF88), II (subunit SDHB), III (subunit UQCRC2), IV (MTCO1) and V (subunit ATP5A), PPARγ coactivator 1 alpha (PGC-1α, Abcam #ab54481, 1:500, rabbit), and MnSOD antibody (Anti-SOD2/MnSOD antibody [2a1], Abcam, #ab16956, 1:500) were obtained from Abcam (Cambridge, MA). Secondary antibodies were IRDye 800RD goat anti-mouse #926-68070 at 1:10,000, IRDye 800RD goat anti-rabbit #926-68071 at 1:10,000 were purchased from LI-COR (Lincoln, NE) and Starbright Blue 700 goat anti-mouse at 1:5,000 #12004159 from Bio-Rad Laboratories (Hercules, California).

### In vivo experiments

The use of animals and protocol were approved by the RMR VA Medical Center IACUC committee. The protocol included guidelines for euthanasia prior to the study end, including weight loss of 10% or more, and behavioral indicators. No adverse events were noted. Animals (male and female Wistar rats, 5 weeks old, n=7-8/group), kept at 2 animals per cage, were housed at either RT (22°C) or TN (29-30°C). Sample size per group was determined based on previous data and experimental design (28). Body temperature was taken superficially and elevated temperature was achieved in those housed at thermoneutral conditions as compared with those housed at room temperature (30.4 ± 0.1°C vs. 27.4 ± 0.1°C, p<0.001, data not shown). Animals were fed a customized diet containing 13% kcal fat (LFD) (Envigo [Teklad]) for 16 weeks. Body weight and food consumption were measured weekly. Endpoint parameters were taken at sacrifice, and all animals were euthanized using isoflurane and exsanguination under anesthetic in the morning following ad libitum food consumption. Female rat estrus cycle was determined according to previously published methodology (29). Briefly, vaginal cells were sampled each morning for three weeks to gauge each animal’s cycle. Cell composition was used to ascertain whether rats were in estrus, proestrus, metestrus, diestrus. Animals were sacrificed while in estrus or proestrus. A randomization program was not used for group assignment, there was no blinding, and cofounders in data acquisition were controlled by altering order of measurements in vivo and ex situ. ARRIVE reporting guidelines were used for these experiments. (Percie du Sert N, et al. The ARRIVE Guidelines 2.0: updated guidelines for reporting animal research.)

### Insulin and glucose intraperitoneal tolerance tests

Insulin tolerance testing (ITT) was done at 1 and 16 weeks of the study. Animals were fasted for 6-hours, and insulin administered by interperitoneal injection of 1U kg-1 body weight. Blood glucose concentrations were taken at 0, 15, 30, 45, 60, and 120 minutes post insulin administration. Glucose tolerance testing (GTT) followed the same protocol, injecting 1.5 g kg-1 body weight of glucose and blood collection done on the same timeline. Baseline concentrations were subtracted in the area under the curve (AUC) analyses.

### Vasoreactivity

Sacrifice of animals occurred at 16 weeks post experimental start. Sacrifices were performed at RT. Aortae and thoracic aortic PVAT were removed from the rats at sacrifice. Vessels with or without PVAT were measured for vasoreactivity using force tension as previously described.(30–33) PVAT was only transplanted to different temperature-housed animals, with sex of the animal being kept the same with PVAT+aorta tissues. Ex situ groups were RT PVAT + RT aorta, TN PVAT + TN aorta, RT PVAT + TN aorta and TN PVAT + RT aorta. For same temperature PVAT and aorta, dissected PVAT was transplanted onto a different section of its aorta to control for manipulation. Transplantation was successful, in part, due to intact adipose connective tissue adhering the PVAT to the aorta. The tissues were adjacent for the whole experiment. To quantify vasoreactivity, tissue (2 mm rings) was mounted on a stainless steel hook and a force-displacement transducer (Grass Instruments Co.) while incubated in a bath at 14.7 mN basal tension; baths contained Krebs buffer (119 mmol L-1 NaCl, 4.7 mmol L-1 KCl, 2.5 mmol L-1 CaCl_2_, 1 mmol L-1 MgCl_2_, 25 mmol L-1 NaHCO_3_, 1.2 mmol L-1 KH_2_PO_4_, and 11 mmol L-1 D-glucose) and continuously bubbled with 95% O_2_ and 5% CO_2_. Constriction was conducted by exposure to 80 mmol L-1 KCl. A phenylephrine dose response curve was also done with doses ranging from 0.002 µmol L-1 to 0.7 µmol L-1. To investigate vasodilation, a dose response curve with ACh was performed with a range of 0.05 µmol L-1 to 20.0 µmol L-1 secondary to phenylephrine constriction. For SNAP response, vessels were pre-constricted with 0.015 µmol L-1 PE to approximate 80% of maximal constriction. SNAP was then added for a final concentration of 2.5 µmol L-1. Data was collected using AcqKnowledge software, and if vessels showed no reactivity to high-potassium, PE, or ACh, they were excluded from analysis. PE constriction was normalized to tissue wet weight, and ACh was normalized as a percentage of relaxation from individual PE maximum constriction.

### Respiration

Mitochondrial respiration was measured using Oroboros Oxygraph-2k (O2k; Oroboros Instruments Corp., Innsbruck, Austria). The aortae (n=8) were placed in a mitochondrial preservation buffer [BIOPS (10 mmol L-1 Ca-EGTA, 0.1 mmol L-1 free calcium, 20 mmol L-1 imidazole, 20 mmol L-1 taurine, 50 mmol L-1 K-MES, 0.5 mmol L-1 DTT, 6.56 mmol L-1 MgCl2, 5.77 mmol L-1 ATP, 15 mmol L-1 phosphocreatine, pH 7.1)], stored on ice, cleaned of fat and connective tissue, and permeabilized by incubation with saponin (40 mg mL-1) in BIOPS on ice on a shaker for 30 minutes. The vessels were then washed for 10 minutes on ice on a shaker in mitochondrial respiration buffer [MiR06 (0.5 mmol L-1 EGTA, 3 mmol L-1 magnesium chloride, 60 mmol L-1 K-lactobionate, 20 mmol L-1 taurine, 10 mmol L-1 potassium phosphate, 20 mmol L-1 HEPES, 110 mmol L-1 sucrose, 1 g L-1 bovine serum albumin, 280 U mL-1 catalase, pH 7.1)]. The vessels were then transferred to MiR06 that had been prewarmed to 37⁰C in the chamber of the O2k. Oxygen concentration in the MiR06 started at approximately 400 nmol ml-1and was maintained above 250 nmol ml-1. Substrates and inhibitors were added to assess respiration rates at several states, including background consumption with carbohydrate or lipid only (state 2), oxidative phosphorylation (+ADP, state 3), maximum oxidative phosphorylation (succinate, state 3S), state 4 (+oligomycin), and uncoupled state (+ FCCP). For the carbohydrate experiment, (pyruvate/malate/glutamate-driven), respiration rates were measured with the final concentrations of 5 mmol L-1 pyruvate + 2 mmol L-1 malate + 10 mmol L-1 glutamate, 2 mmol L-1 adenosine diphosphate (ADP), 6 mmol L-1 succinate, 4 mg mL-1 oligomycin, and 0.5 mmol L-1 stepwise titration of 1 mmol L-1 carbonyl cyanide 4-trifluoromethoxy) phenylhydrazone (FCCP) until maximal uncoupling (uncoupled state). In the lipid experiment (palmitoylcarnitine-driven respiration), rates were measured with 5 µmol L-1 palmitoylcarnitine + 1 mmol L-1 malate, 2 mmol L-1 ADP, 2 mmol L-1 glutamate + succinate, 4 mg mL-1 oligomycin, and 1 mmol L-1 stepwise titration of FCCP. Cytochrome c (10 mmol L-1) was used to determine mitochondrial membrane integrity. Vessels were dried overnight at 60°C and weighed for dry weight normalization. Using the Oroboros DatLab software, traces were analyzed by averaging a stable trace segment of 3-5 minutes for each state.

### Western blotting

Aortae and aortic fat samples were flash-frozen in liquid nitrogen and later processed in mammalian lysis buffer (MPER with 150 mmol L-1 sodium chloride, 1 mmol L-1 of EDTA, 1 mmol L-1 EGTA, 5 mmol L-1 sodium pyrophosphate, 1 mmol L-1 sodium orthovanadate, 20 mmol L-1 sodium fluoride, 500 nmol L-1 okadaic acid, 1% protease inhibitor cocktail). Aortae and aortic fat were ground under nitrogen with a mortar and pestle, and homogenized at 4°C and centrifuged first at 1,000 x g for 2 min, and supernatants subsequently centrifuged 16,400 x g at 4°C for 10 min. A Bradford protein assay was used to measure the protein concentration of the lysates. Protein samples (15 µg to 40 µg) in Laemmli sample buffer (LSB, boiled with 100 mmol L-1 dithiothreitol [DTT]) were run on precast SDS-4-15% polyacrylamide gels. Proteins were transferred to PVDF membranes. Quantity One, Bio-Rad, was used to evaluate protein loading. Blots were probed with antibodies described above and left overnight at 4°C. Fluorescent secondary antibodies were applied following the primary antibody incubation (1:10,000 IRDye800CW, 1:10,000 IRDye680RD and Starbright Blue 700), 1 hour at room temperature, protected from light). Total protein was measured using the activation of the stain-free gels, according to manufacturer instructions for the ChemiDoc on the ChemiDoc Imaging System (Bio-Rad, Hercules, CA) using the Quantity One 1-D Analysis software (Bio-Rad, Hercules, CA). All protein data has been normalized to loading control and total protein expression by quantifying the total protein in the entire lane. Blots are imaged first for total protein, then exposed to specific wavelengths corresponding to the protein target of interest, to ensure normalization per each target and each associated process such as washing. For determining the ratio of phosphorylated signal to total signal, antibodies were probed on the same blot using different animal primary antibodies between the phosphorylated and total protein (rabbit vs. mouse) allowing for two color detection and analysis when used with secondary florescent antibodies with differing wavelengths (IRDye 680RD and IRDye 800CW).

### Hematoxylin & eosin (H&E) and immunohistochemistry (IHC) tissue staining

Aortic fat samples were collected from sacrificed animals. These tissue samples were fixed in 10% formalin and embedded in paraffin using a Leica HistoCore Pearl tissue processor and Tissue Tek III embedder. Each tissue mold was processed into 5μm sections using a Leica RM2235 Microtome. To prepare for staining, each slide was deparaffinized and rehydrated. Tissue sections analyzed by H&E were washed twice in 1xPBS for 2 minutes each, incubated in Hematoxylin for 30 seconds, and incubated in the following: 0.1% ammonia for 10 seconds, 95% ethanol for 2 minutes, eosin for 2 minutes, acidified 70% ethanol for 2 minutes, acidified 95% ethanol for 1 minute, 100% ethanol for 2 minutes, and CitriSolv hybrid twice for 2 minutes each. BAT and WAT morphology was determined by analyzing lipid droplet size and multilocular appearance of adipose tissue in brightfield. BAT phenotype was also observed as distinctly multilocular. PVAT samples contained both BAT and WAT morphology, and amount of each adipocyte type was quantified as a ratio and normalized to total tissue area using Keyence BZ-X800 software. For IHC, each slide was taken post hydration for antigen retrieval by incubation in DivaDecloaker solution in a pressure cooker for 10 minutes. UCP-1 polyclonal antibody diluted 1:125 in 1% Goat Serum was then added to each slide and incubated for 1 hour. Cy5 goat anti-rabbit antibody was diluted 1:500 in 1% goat serum and added to each slide for 10 minutes. Conjugated wheatgerm agglutinin diluted 1:5 in 1% goat serum was added to the solution on each slide and incubated for another 10 minutes. Mounting media including DAPI was also used. Each tissue section was imaged 20x with exposures of 1/35 seconds for DAPI and 1/2.5 seconds for Cy5 (UCP-1) using a Keyence BZ-X800 fluorescent microscope. Analysis was done using Keyence BZ-X800 analyzing software. This software quantifies fluorescent images, including DAPI staining, and can be used to obtain total fluorescent signal. UCP-1 total signal was normalized to total tissue area.

### Adiponectin ELISA

Adiponectin levels were measured in PVAT samples using a Quantikine ELISA Rat Total Adiponectin/Acrp30 kit (R&D systems, Cat.#RRP300). Flash frozen samples of PVAT were made into lysate preparations and measured for protein concentrations using a Bradford assay. Samples were then analyzed for adiponectin concentrations per the manufacturers protocol and normalized to amount of tissue sample used (mg).

### Mechanical Measurements of Aortic Stiffness

Measurements were done based on a previous protocol (34). Approximately 2mm long PVAT + aortic tissue samples were freshly collected at sacrifice, added to chilled 1xPBS and stored in -80°C until analysis. Axial bisection of the arteries produced circumferential rectangular tissue segments for mechanical testing. Measurements were taken using digital images and calipers. Using known measurements taken with the calipers, vessel thickness, vessel length, and gage length were calculated using MATLAB software, providing a baseline set of measurements for each aorta. Tissue stress-strain measurements were acquired using an MTS, Insight 2 (MTS Systems, Eden Prairie, MN) material testing system, with ancillary environmental chamber and 2-N load cell. Testing was conducted in 1xPBS heated to 37°C in an attached water bath. Tissues were preconditioned by 5 full extensions-relaxation cycles before data collection. The strain rate for all mechanical tests was 10% strain per second. Tissues were stretched in repetitions of 5, in increasing increments of 5% strain, starting at 50%, until the collagen-dependent, strain-stiffening response became fully developed, typically around 60-80% strain. Data analysis was conducted using custom-written software provided by Kendall Hunter at University of Colorado (Matlab R2022a, The Math Works, Natick, MA).

### Statistical analysis

To analyze data with time/dose, such as PE or ACh, along with sex and temperature, we employed a repeated measures ANOVA, along with a mixed-effects model. For data without a time or dose component, we employed a two-way ANOVA for the variable temperature and sex. Tukey’s post-hoc analyses were conducted within ANOVA tests. A p value of less than 0.05 was used as the cutoff for statistical significance in all tests. A p value of equal or less than 0.08 was considered indicative of data trends approaching significance. Symbols denoting significance are consistent between Figures 1 and 2A-D to denote repeated measures ANOVA for dosage. Symbols used in all other figures reflect post-sacrifice analysis, without any repeated measures.

**Figure 1A and B:**
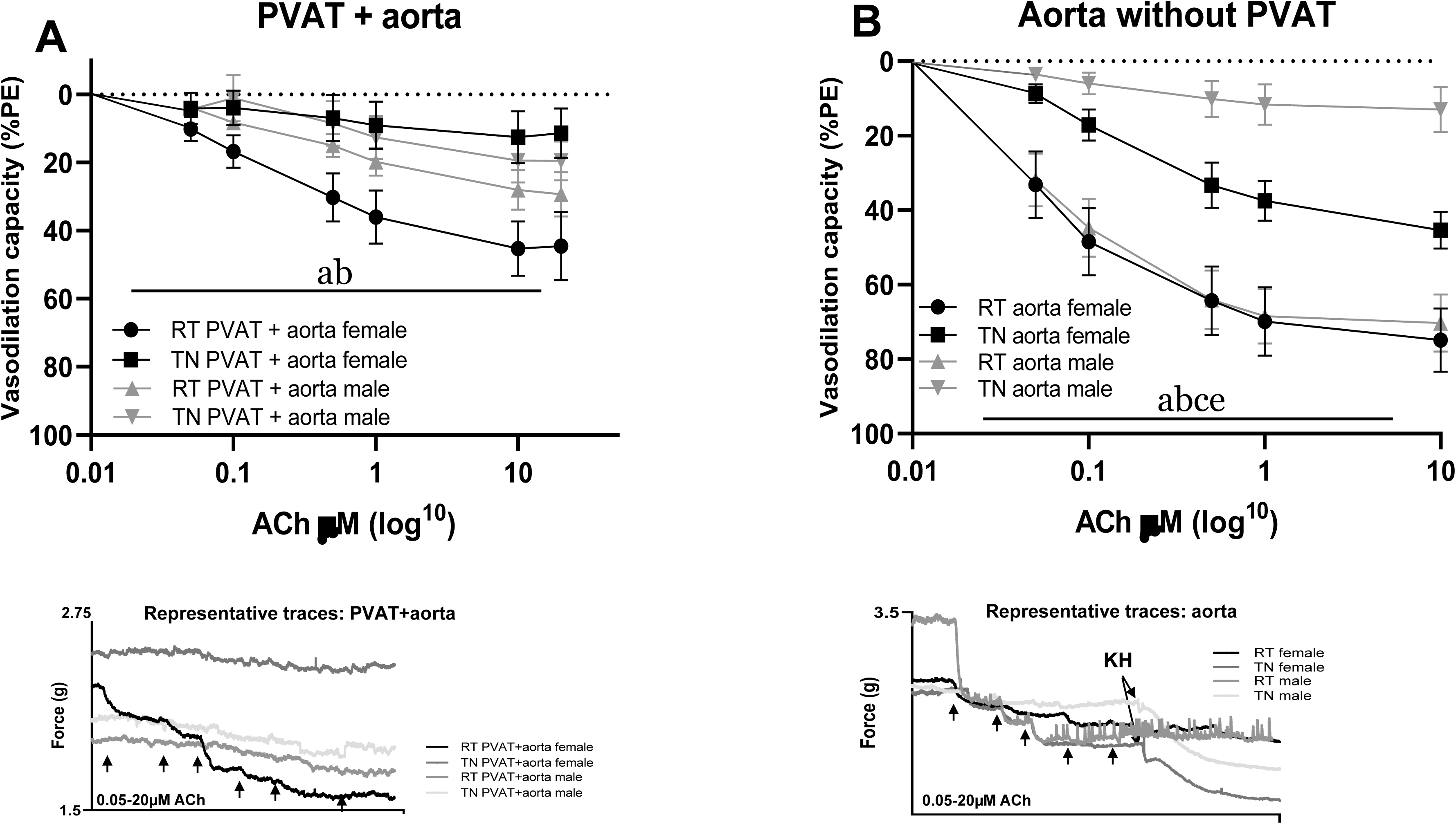
Vasoreactivity of aortae in response to acetylcholine (ACh, Figure 1A) with PVAT or cleaned of PVAT. Aortae with PVAT (A) or cleaned, intact vessels (B) were attached to a force transducer and exposed to an increased dose of Ach, n=7-8. ACh dose response (µM log^10^) is expressed as a percentage of fully PE-constricted vessels in all rats. Effects of temperature (^a^p<0.05), ACh x temperature interaction (^b^p<0.05), ACh x sex interaction (^c^p<0.05), all variables interaction (^d^p<0.05), mixed-effects and/or repeated measures three-way ANOVA, Figures 1A and B. Data are mean ± SEM. Representative traces of each graph are shown below.

**Figure 2A-F:**
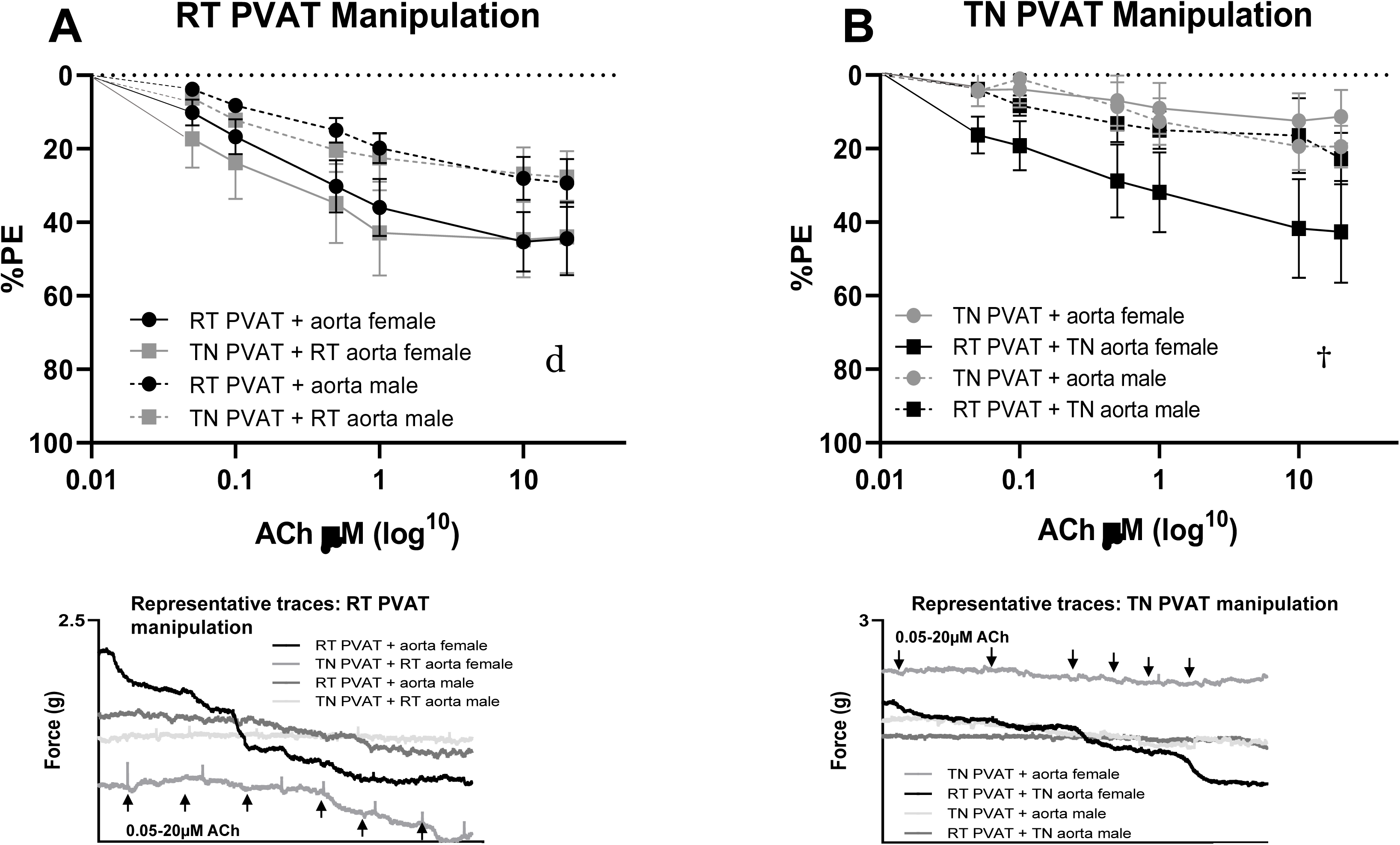
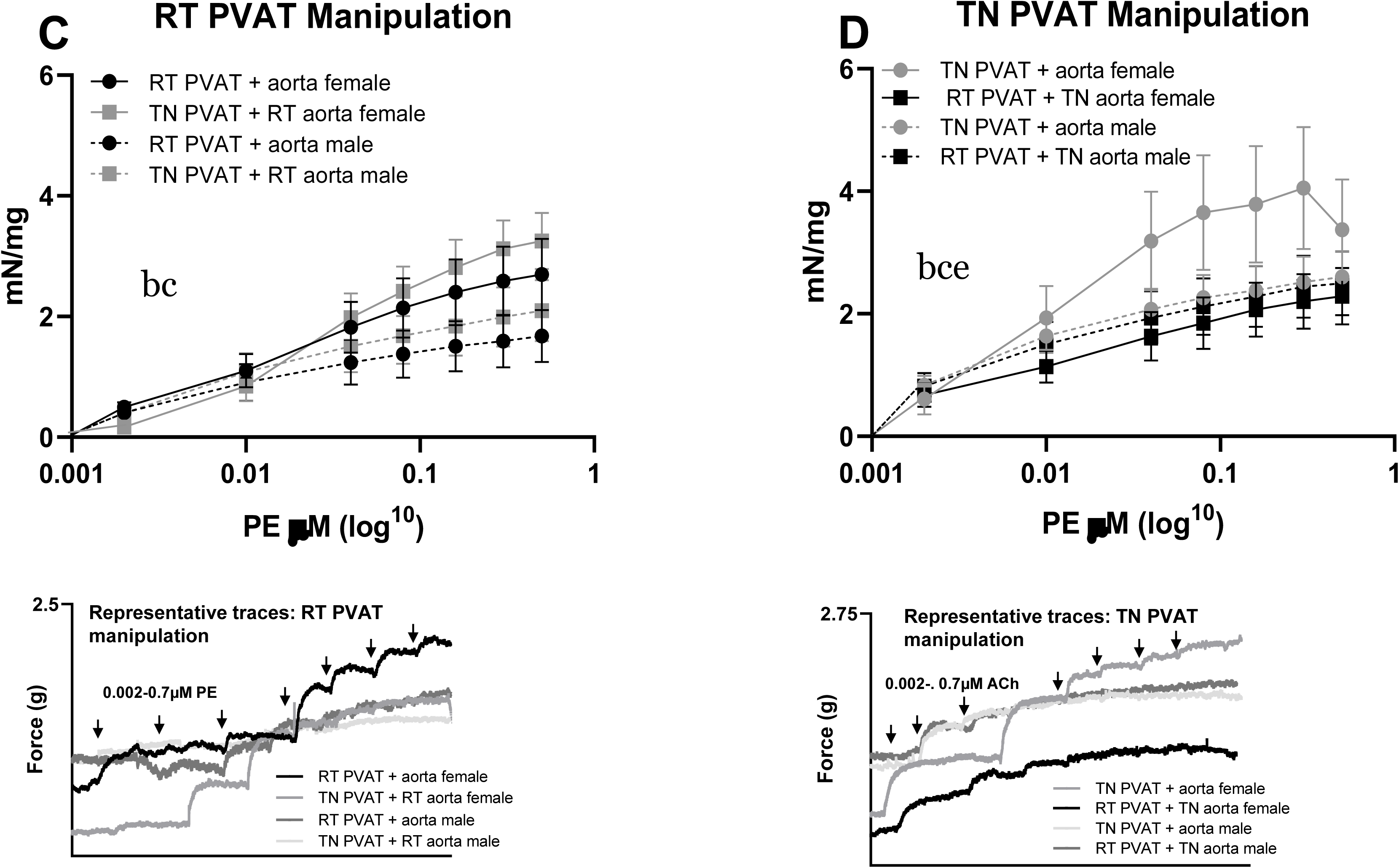
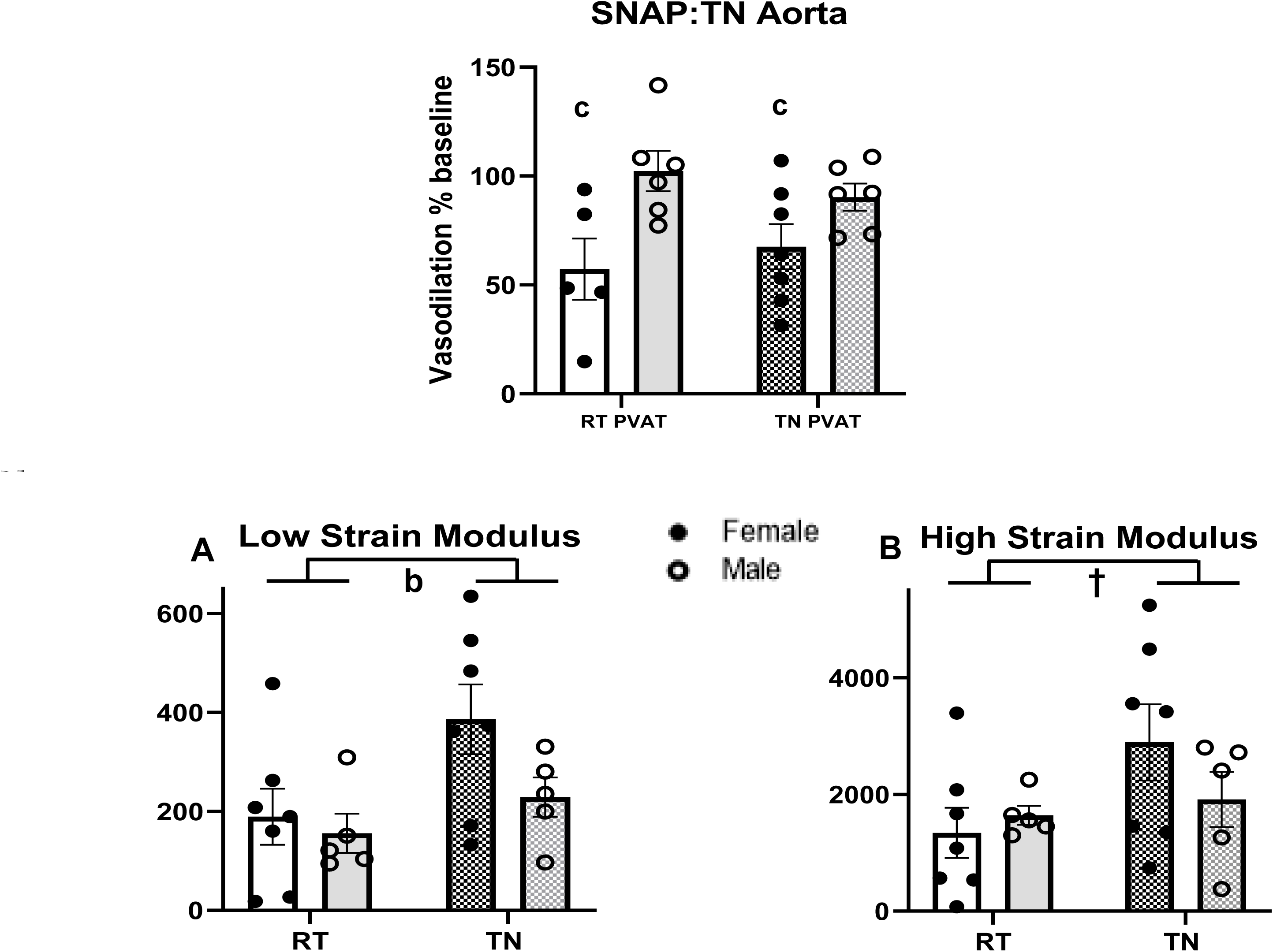
Vasoreactivity of PVAT + aorta in response to acetylcholine (ACh, Figure 2A and B) or phenylephrine (PE, Figure 2C and D). PVAT + aorta, either controls (RT PVAT + RT aorta or TN PVAT + aorta) or manipulations (RT PVAT + TN aorta or TN PVAT + RT aorta) were attached to a force transducer and exposed to an increased dose of either ACh (A and B) or PE (C and D), n=7-8. Controls used in Figures 1A and B are also depicted here. ACh dose response (µM log^10^) is expressed as a percentage of fully PE-constricted vessels (A). PE dose response is expressed as mN/mg normalized to vessel wet weight (B). Effects of temperature (^a^p<0.05), ACh/PE x temperature interaction (^b^p<0.05), ACh/PE x sex interaction (^c^p<0.05), sex (^d^p<0.05), all variables interaction (^e^p<0.05, ^†^p<0.08), mixed-effects and/or repeated measures three-way ANOVA, Figures 1A and B. Data are mean ± SEM. Representative traces of each graph are shown below. **Aortic response to SNAP (E) and stiffness measured via strain modulus (F).** Aortae with PVAT manipulation exposed to SNAP and measured for dilation as percentage of pre-constriction (E). Aortae together with PVAT was measured for elastin and collagen contributions to stiffness using low (A) and high (B) strain modulus, n=7-8. Interaction effect ^a^p<0.05, effect of temperature ^b^p<0.05, effect of sex ^c^<0.05, interaction effect ^†^p=0.08, two-way ANOVA. Data are mean ± SEM.

## Results

### Housing temperature impacted body weight and insulin sensitivity

Significant temperature effects were noted in weights (p<0.01, Table 1A), with rats housed at TN showing lower weights than those housed at RT (p<0.01, Table 1A). There was a significant sex effect, with male rats weighing more than females (p<0.01, Table 1A). There was a significant effect of sex on glucose concentrations (p<0.01, Table 1A), with male rats having higher glucose concentrations than females (p<0.01, Table 1A). There were significant sex effects on weight of gonadal-epidydimal/ovarian fat (p<0.05, Table 1B). During a GTT, no significance differences were noted, however males had significantly less insulin sensitivity after 16 weeks at TN housing (Table 1B, p<0.05). This change in insulin sensitivity was not observed in the female rats.

**Table 1A.**
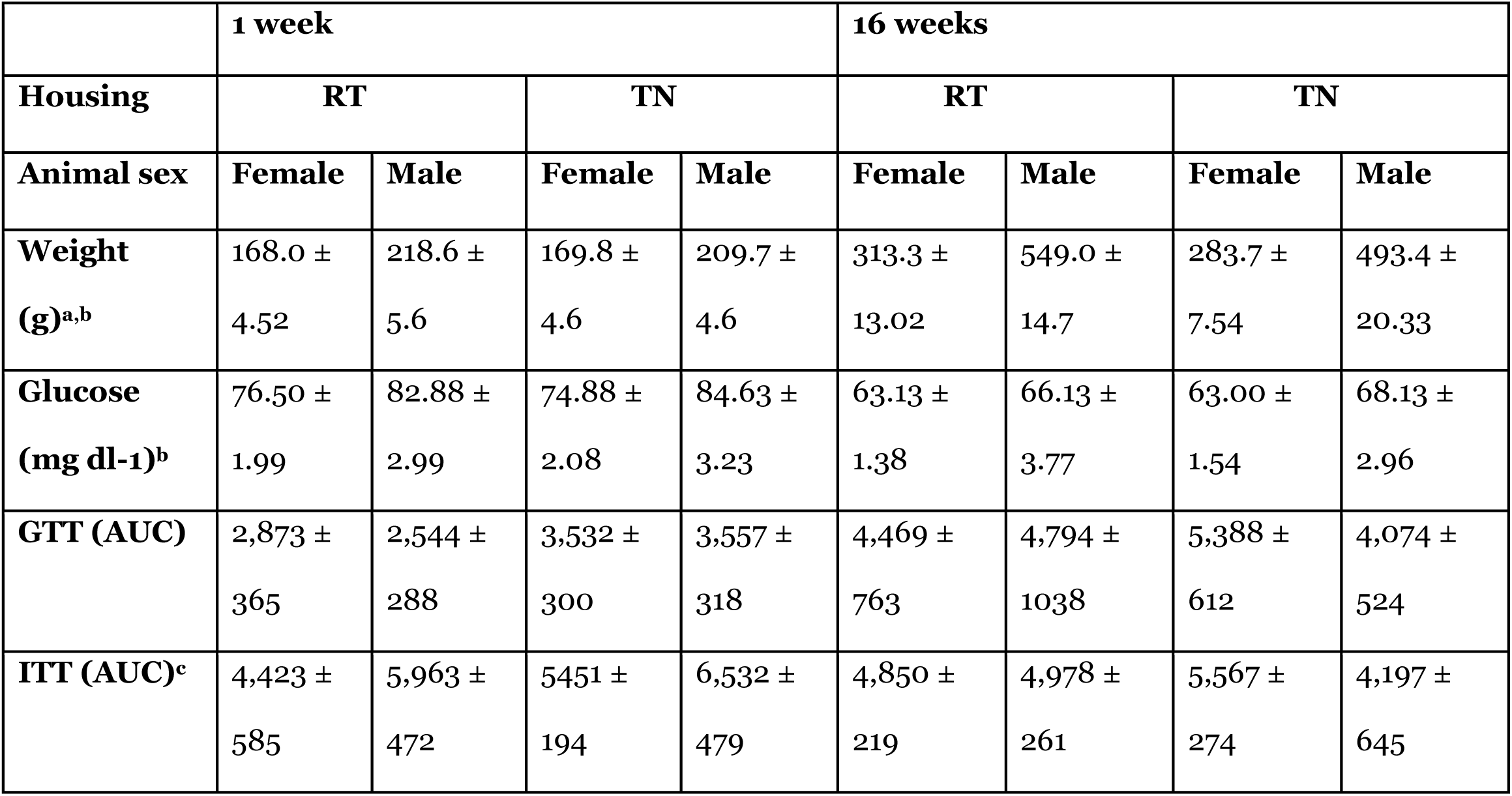
Animal weight, fasting glucose concentrations, and area under the curve (AUC) for glucose and insulin tolerance testing (GTT, ITT) at 1 and 16 weeks of treatment. Animal biological parameters were assessed at 1 and 16 weeks, n=7-8. GTT and ITT were conducted in fasted animals, with AUC analyzed. ^a^p<0.05 temperature, ^b^sex, ^c^time x sex, three-way ANOVA, mean ± SEM.

**Table 1B.** Animal adipose depots at 16 weeks of treatment. Gonadal-epididymal (G-E) or ovarian (O), along with cardiac, adipose depots were measured at sacrifice, n=7-8. Analysis was conducted on adipose depot weight along as well as weight normalized to body weight. ^a^p<0.05 temperature, ^b^sex, ^c^time x sex, three-way ANOVA, mean ± SEM.

|  | **16 weeks** |  |  |  |
| --- | --- | --- | --- | --- |
| **Housing** | **RT** |  | **TN** |  |
| **Animal sex** | **Female** | **Male** | **Female** | **Male** |
| **Cardiac fat (mg)<sup>b</sup>** | 117.01 ± 19.03 | 223.93 ± 27.41 | 113.01 ± 19.34 | 234.14 ± 29.88 |
| **Cardiac fat/body weight (mg/g)** | 0.38 ± 0.06 | 0.41 ± 0.05 | 0.40 ± 0.06 | 0.46 ± 0.04 |
| **G-E/O fat (g)<sup>b</sup>** | 4.22 ± 0.22 | 12.18 ± 1.03 | 3.15 ± 0.39 | 10.72 ± 1.05 |
| **G-E/O fat/body weight (mg/g)<sup>b</sup>** | 13.29 ± 0.98 | 22.04 ± 1.56 | 11.22 ± 1.54 | 18.47 ± 2.91 |

### PVAT impacts aortic vasoreactivity in a sex-dependent manner

Aortae together with PVAT from animals housed at TN or RT for 16 weeks, as well as aortae cleaned of PVAT, were assessed for dilation response to vasodilator ACh (Figure 1A and B). There was a significant effect of temperature exposure on aortae response to vasodilator ACh, resulting in greater vasodilatation in RT-housed rat aortae (p<0.05, Figure 1A). When analyzed for sex differences alone, only TN females showed a significantly diminished vasodilation response (p<0.05, Figure 1A). Aortae with PVAT removed showed a significantly lesser response from animals housed at TN (p<0.05, Figure 1B). Both male and female rats had significantly diminished vasodilation response at TN (p<0.05, Figure 1B).

### RT PVAT restored vasodilation in TN aortae in a sex-dependent manner

Aorta together with its own PVAT, or aorta with PVAT from an oppositely-housed animal, were analyzed in response to vasodilator ACh and vasoconstrictor PE (Figure 2A-D). In rat aorta from RT-housed rats, only sex differences in response were significant (p<0.05 for both, Figure 2A and 2C). However, RT PVAT significantly restored both vasodilation and vasoconstriction response in TN aorta (p<0.05 for both, Figure 2B and 2D); this improvement was significant in females only (p<0.05 for both, Figure 2B and 2D). To determine the contribution of the endothelium, we exposed PVAT with aortae to nitric oxide donor SNAP. With TN aorta with either TN or RT PVAT, there was a significant effect of sex with female aortae showing less vasodilation than males (p<0.01, Figure 2E).

### Housing temperature increased both elastin and collagen-related aortic stiffness

We used a strain modulus to gauge aortic stiffness in animals housed at either RT or TN (Figure 2E and F). We observed a significant effect of temperature on the contribution of elastin (low strain modulus) in stiffness (p<0.05, Figure 2E), showing the most elevated stiffness in TN-housed females. In gauging collagen’s role in stiffness (high strain modulus), TN-housed animals had a higher collagen contribution to stiffness (Figure 2F), with TN-housed females showing the highest.

### TN promoted whitening of PVAT

We characterized PVAT from all animals using phenotypic and molecular markers of BAT and WAT morphology, as illustrated in Figure 3. In microscopic analysis of H&E stained PVAT, multilocular phenotype, characteristic of BAT, was significantly diminished in rats housed at TN (p<0.01, Figure 3). The percent of the total area of UCP-1 using florescence microscopy on PVAT samples was non-significantly decreased in all animals housed at TN(p=0.06, Figure 3), with females showing the most precipitous decrease. There was a significant interaction of sex and temperature with BAT phenotypic regulator PRDM16 expression (p<0.05, Figure 3), with significant differences between the sexes (p<0.05, Figure 3). Male PVAT aortae PRDM16 expression decreased at TN (Figure 3). Adiponectin concentrations in PVAT were significantly less in PVAT of all animals housed at TN (p<0.05, Figure 3). Female PVAT showed a larger decline of adiponectin than males (Figure 3).

**Figure 3A and B:**
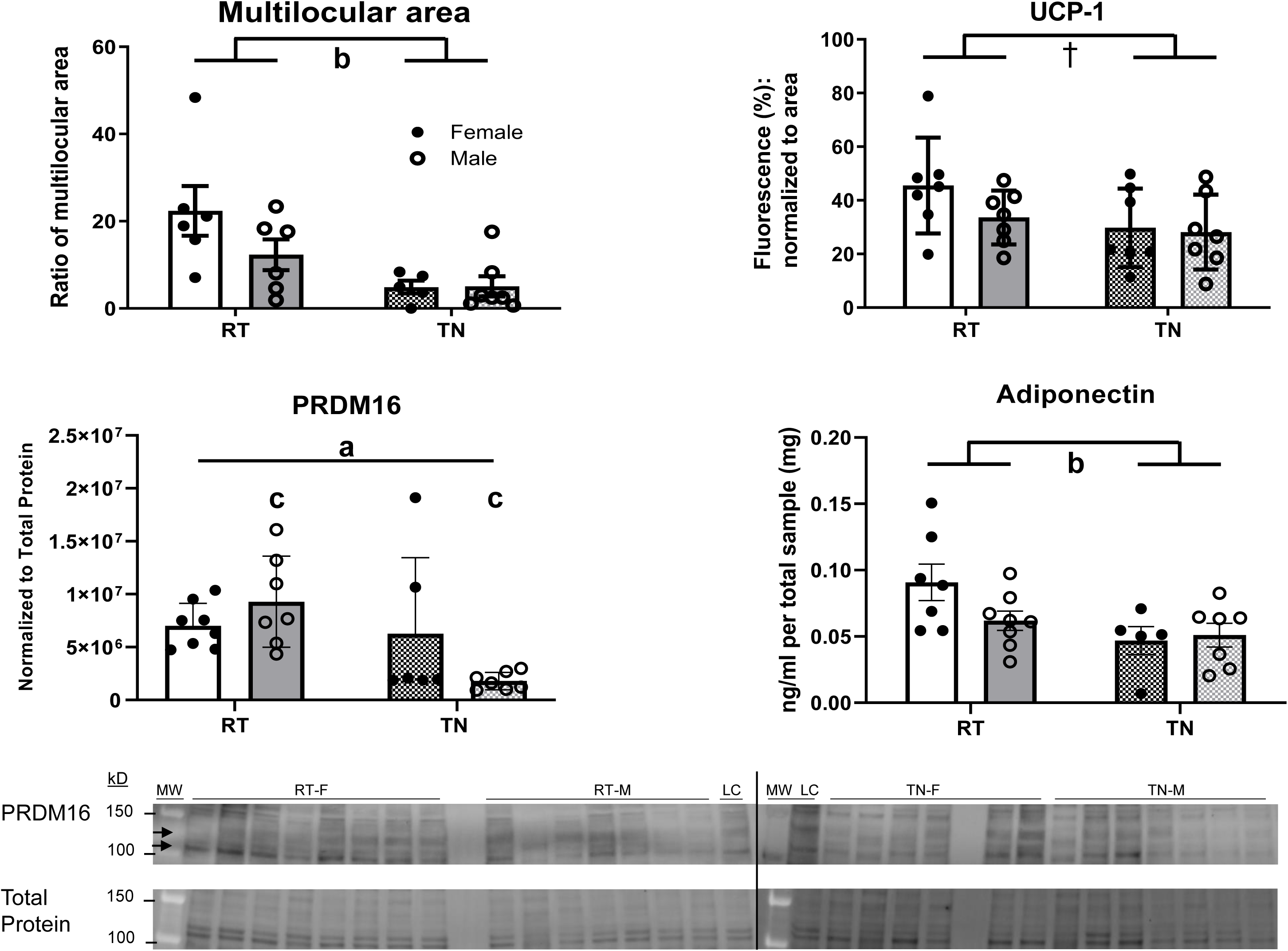
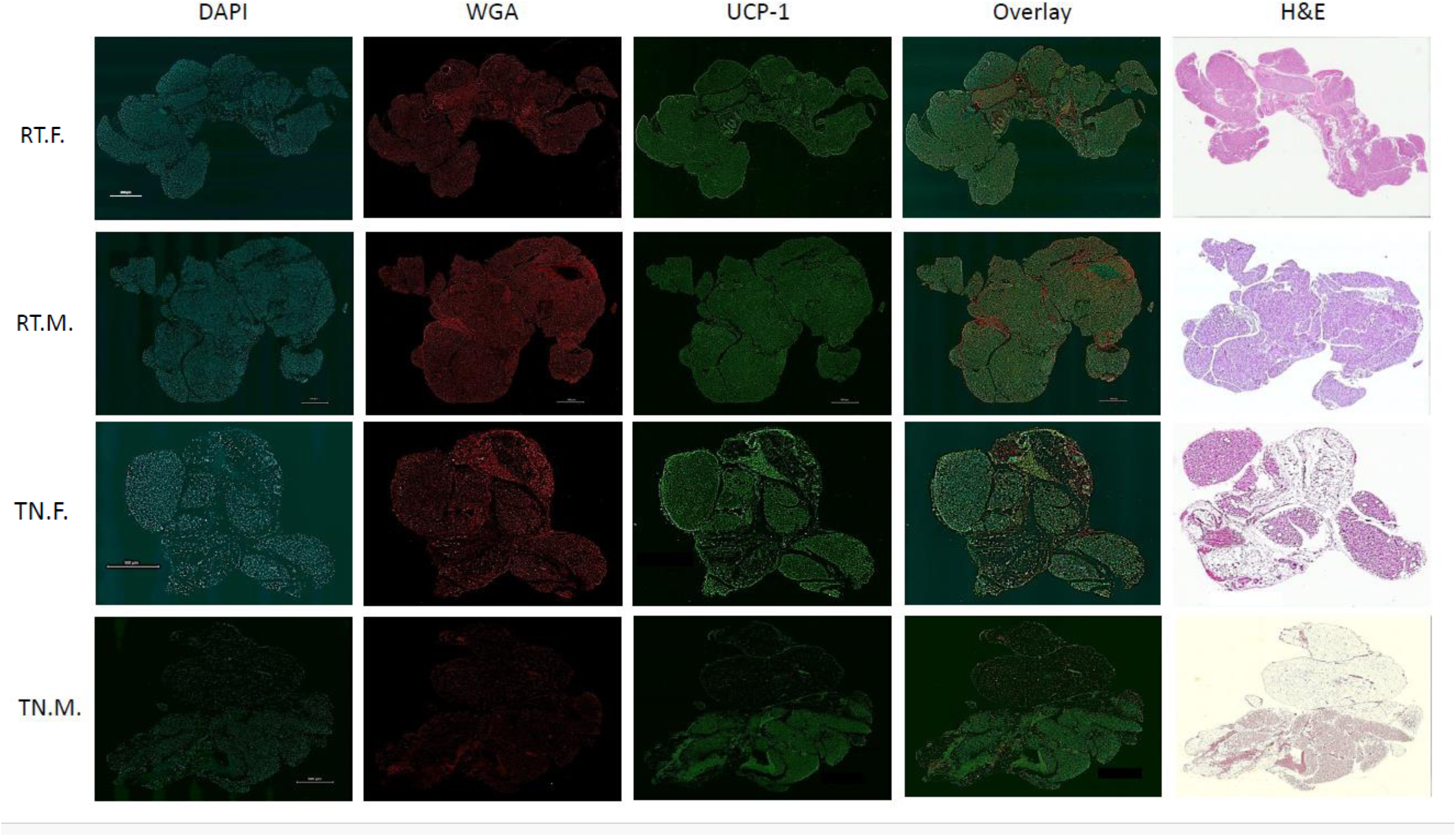
Characterization of PVAT phenotype (A) and representative images (B). BAT or WAT phenotype of PVAT was characterized by adipocyte characteristics and multilocular phenotype using H&E or florescent staining and microscopy analysis to determine multilocular area and amount of UCP1, n=7-8.. For immunofluorescence, PVAT was stained with DAPI (nucleus), Wheat germ agglutinin (actin), and UCP-1. The percentage of the UCP-1 area was normalized to the total area of the tissue. PRDM16 was measured using Western blotting and ELISA was used for adiponectin concentrations. Interaction effect ^a^p<0.05, effect of temperature ^b^p<0.05, effect of sex ^c^<0.05, interaction or temperature effect ^†^p=0.08, two-way ANOVA. Data are mean ± SEM.

### Endothelial and mitochondrial regulatory cellular signaling was altered in TN-housed animals

Total expression of PVAT eNOS was significantly dampened at TN (p<0.05, Figure 4A), with significant elevation of PVAT peNOS and peNOS:eNOS at TN (p<0.05 for both, Figure 4A). PVAT expression of PGC-1α was not significantly different between groups or sexes (p<0.05, Figure 5A). Complexes I, II, and IV were all dampened with TN housing (p<0.05 for complex I and II, p<0.08 for complex IV, Figure 5A), and complex III was significantly elevated in those housed at TN (p<0.05, Figure 5A); males’ complex V expression was significantly elevated at both RT and TN (p<0.05, Figure 5A). Phosphorylated eNOS expression and peNOS:eNOS ratio were significantly less in aortae those housed at TN (p<0.05 for both, Figure 4B). There was a significant interaction of temperature and sex on total eNOS expression, resulting in higher expression of eNOS in aortae from male rats housed at TN (p<0.05, Figure 4B). Upstream of mitochondrial function, aortae PGC-1α expression was significantly higher in male aorta (p<0.05, Figure 5B). Mitochondrial oxphos complexes I and IV were significantly dampened in aortae from TN-housed animals in comparison to those housed at RT (p<0.05 for both, Figure 5B). Complexes II, III and V were expressed significantly higher in male aorta than females (p<0.05 for all, Figure 5B).

**Figure 4A and B.**
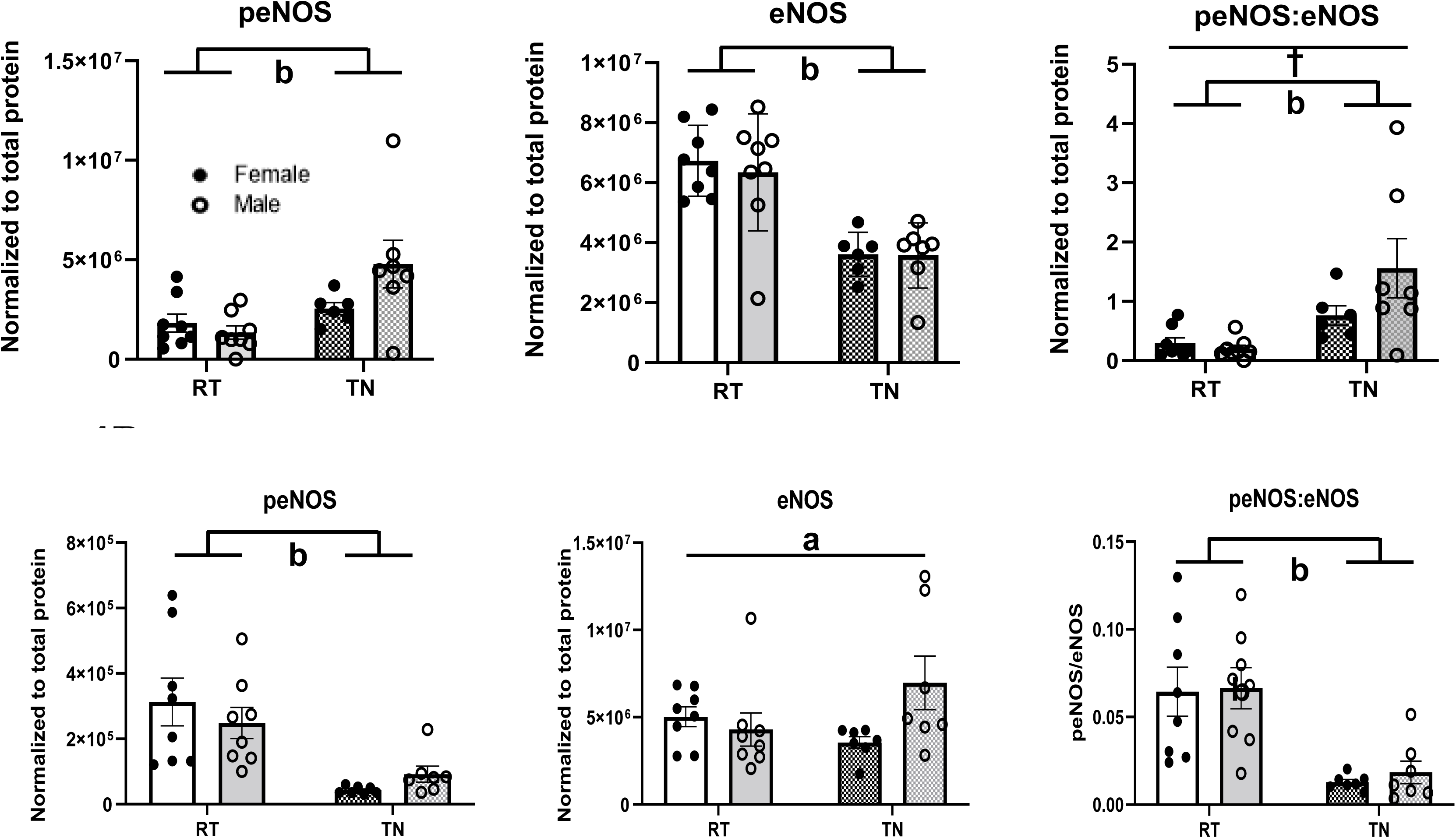
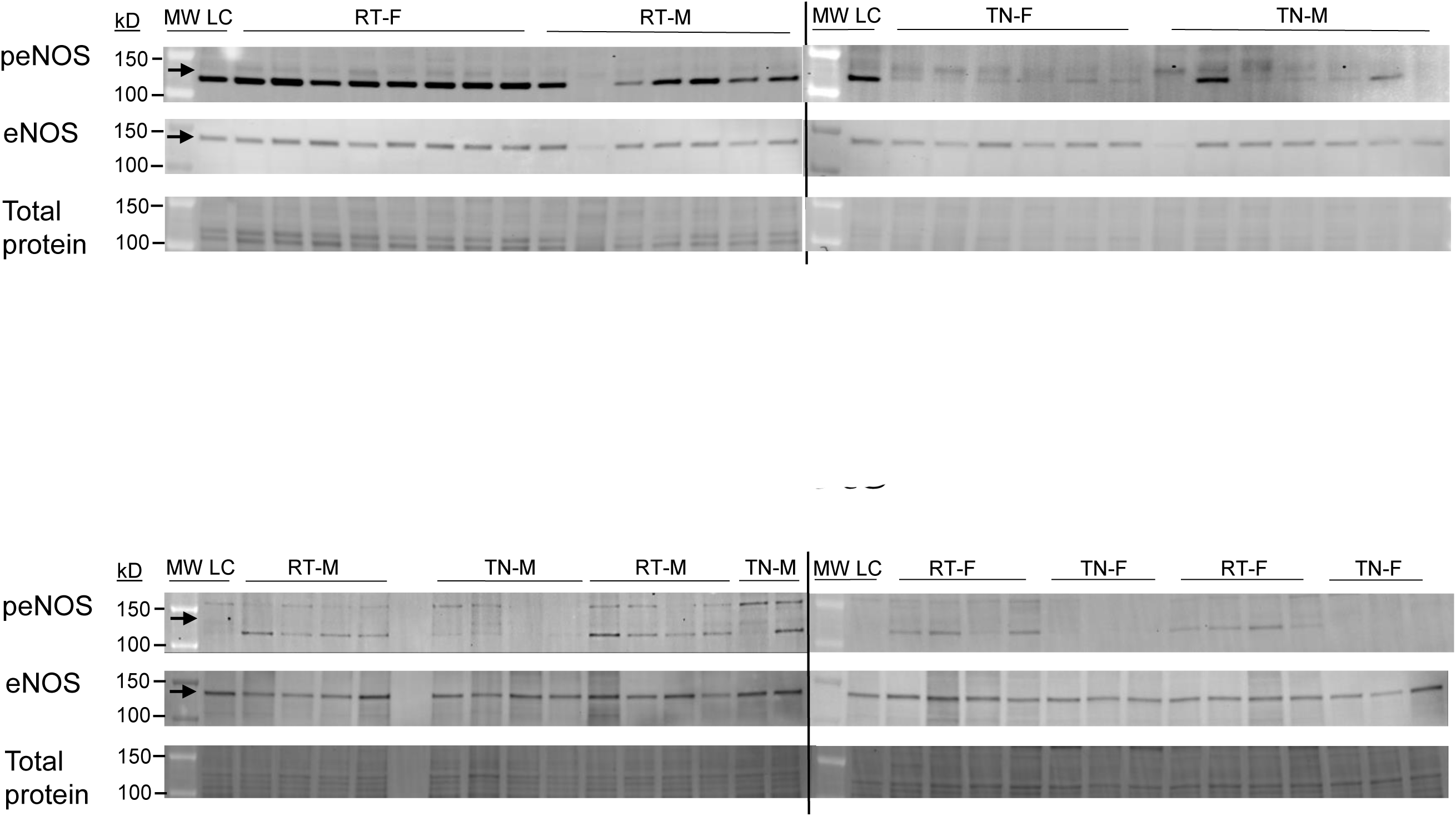
Protein expression of eNOS in PVAT and aortae. Aorta and PVAT tissue was processed for protein analysis via Western blot analysis, including specific activity, n=7-8. Blots were probed for peNOS, and eNOS. Interaction effect ^a^p<0.05, effect of temperature ^b^p<0.05, effect of sex ^c^<0.05, effect of sex or temperature ^†^p=0.08, two-way ANOVA. Data are mean ± SEM.

**Figure 5A and B.**
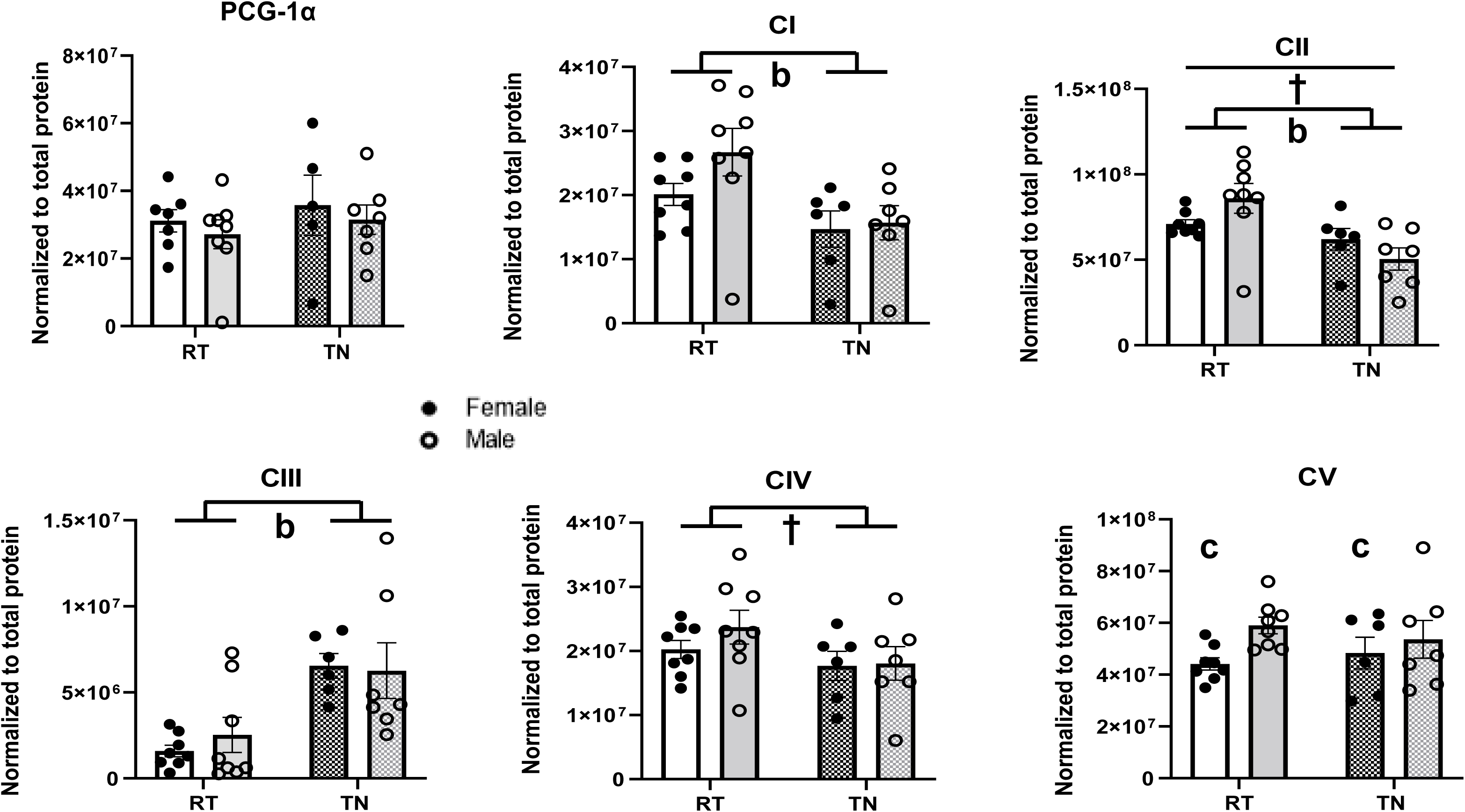
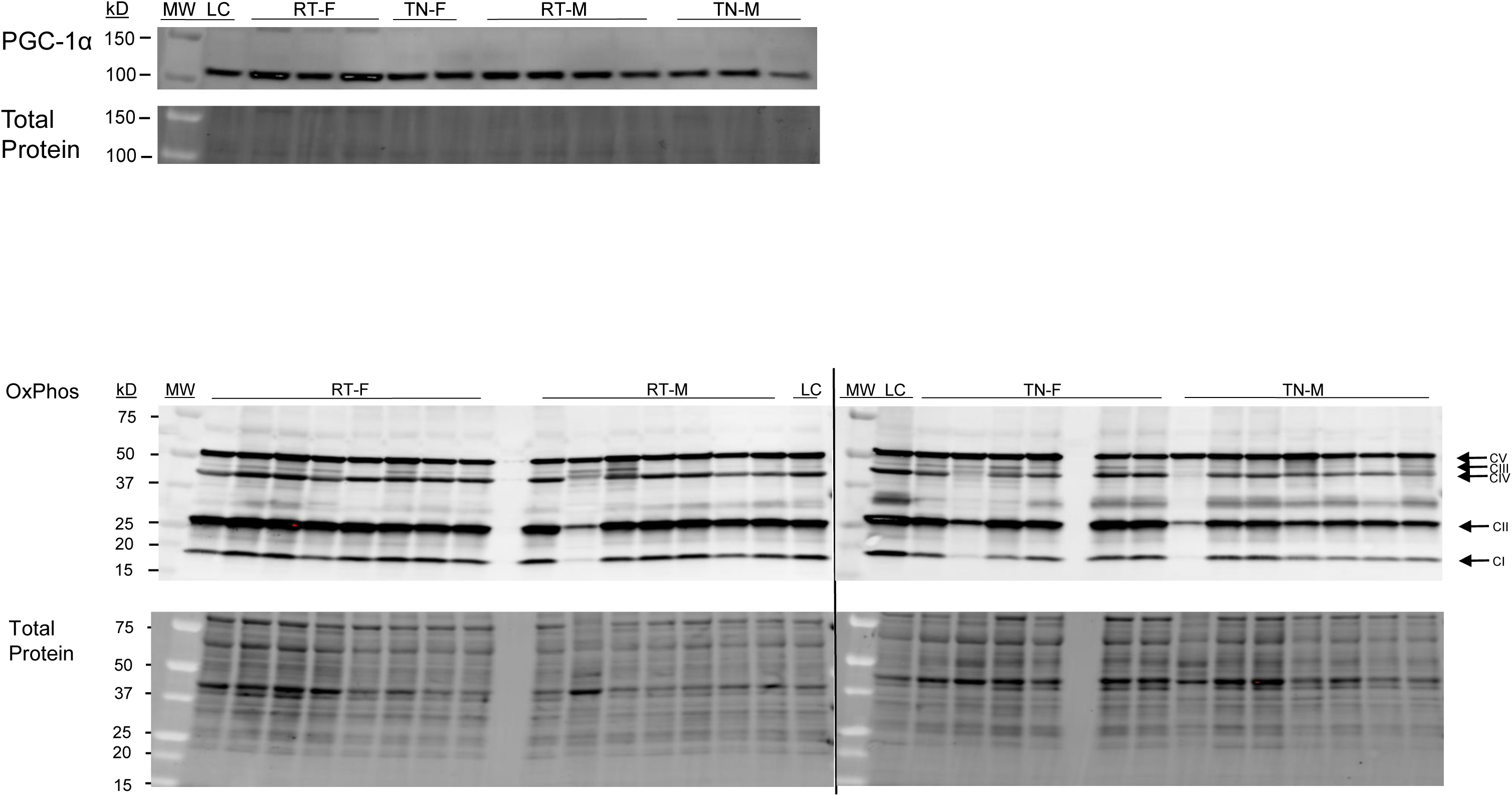
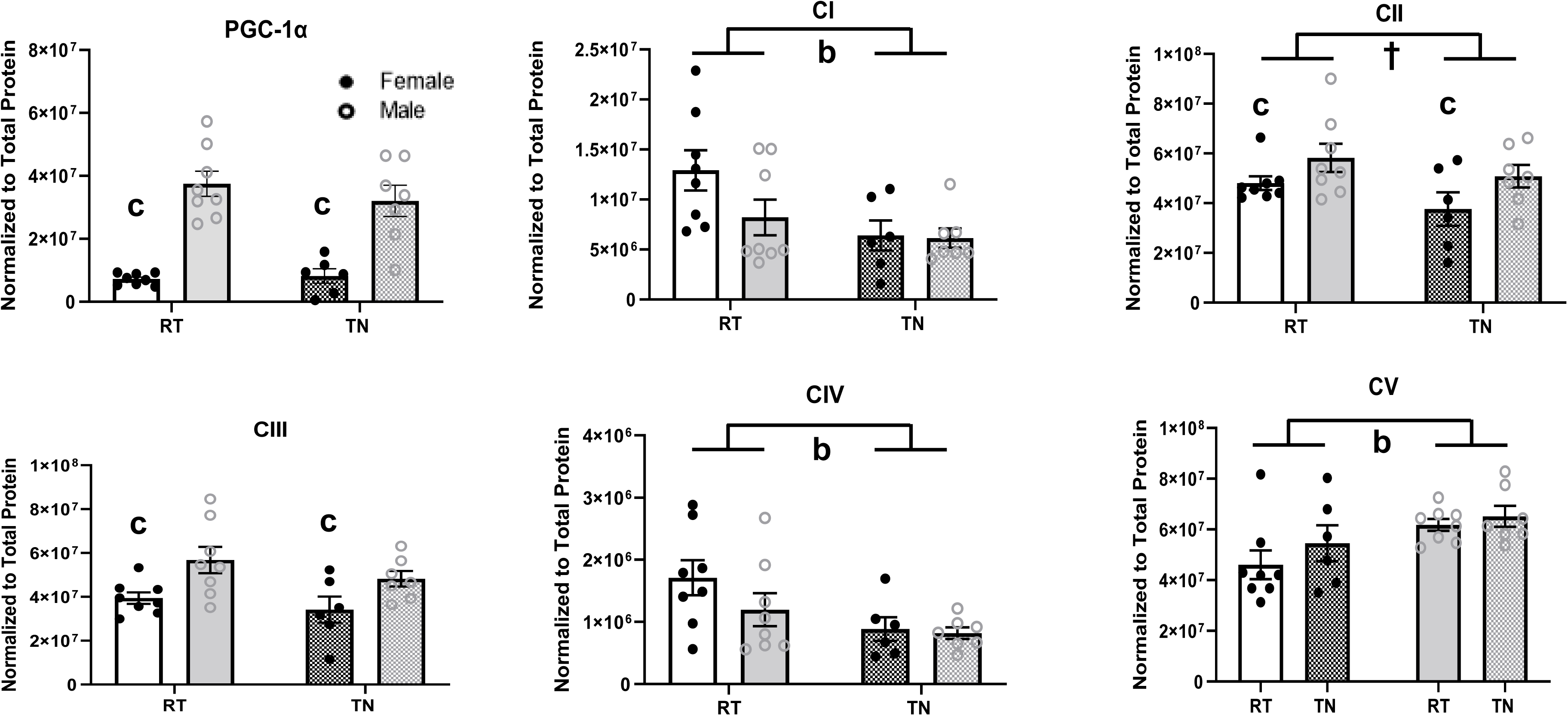
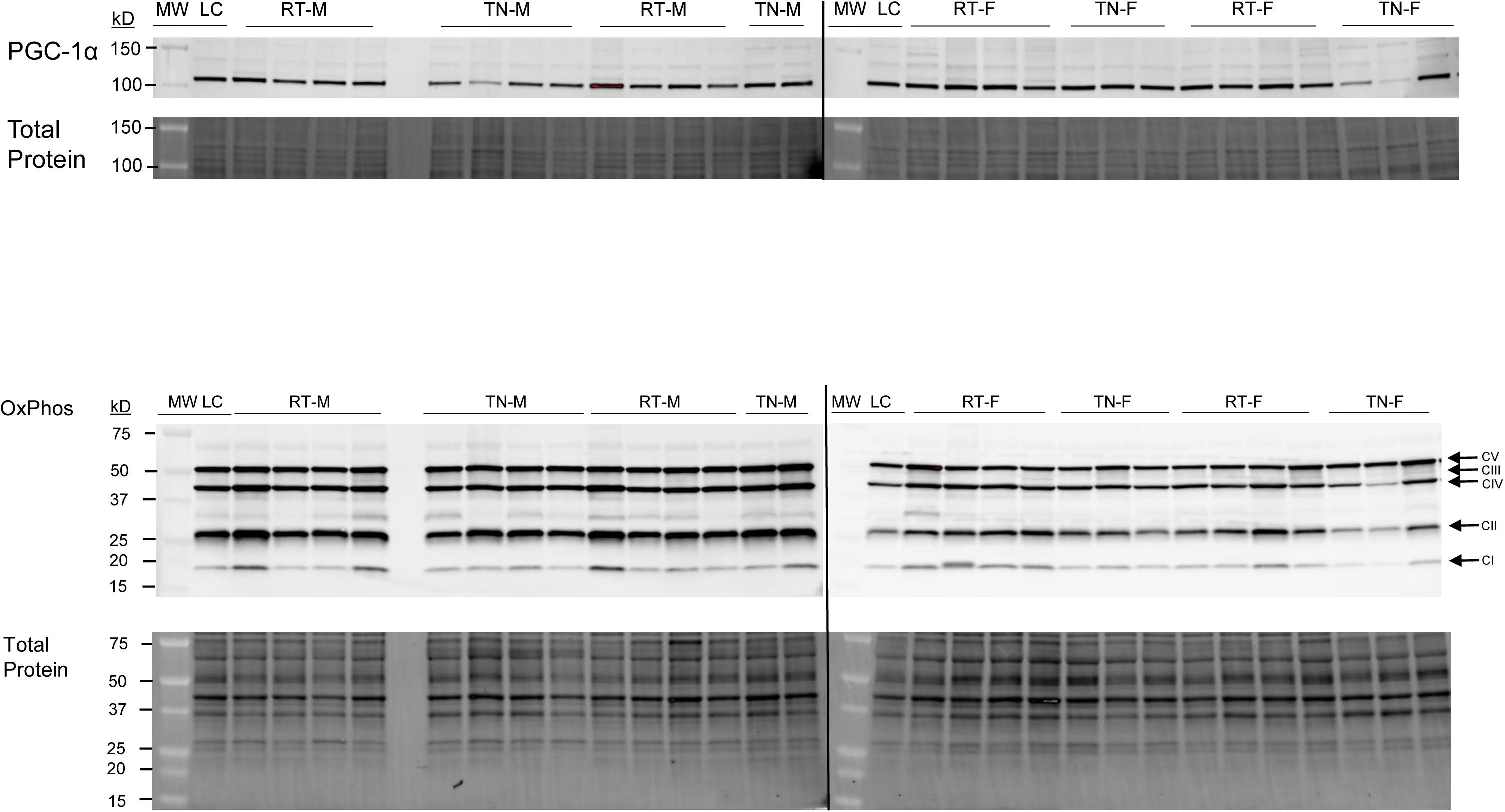
PVAT and aortae protein expression of PCG-1α and mitochondrial complexes. Aortae and PVAT tissue was processed for protein analysis via Western blot analysis, including PCG-1α and mitochondrial complexes, n=7-8. Blots were probed for PCG-1α and mitochondrial complexes. Interaction effect ^a^p<0.05, effect of temperature ^b^p<0.05, effect of sex ^c^<0.05, ^†^p=0.08, 2-way ANOVA, two-way ANOVA. Data are mean ± SEM.

### TN housing resulted in lower mitochondrial respiration in the context of lipid substrates

In multiple states representing ATP-coupled respiration (state 3S), membrane potential maintenance (state 4) and uncoupled maximal respiration, aortae from rats housed at TN showed significantly less respiration rates compared with those of RT house rat aorta (p<0.05 for all, Table 2). Aorta from TN-housed rats also had significantly elevated respiratory control ratio, an indicator of oxphos activation versus reserve (RCR, p<0.05, Table 2). In experiments with carbohydrate experiments, TN-housed aortae also showed significantly less state 4 respiration than those from RT-housed rats (p<0.05, Supplemental Table 1). State 3S respiration was also lower (Supplemental Table 1).

**Table 2.** Mitochondrial respiration, lipid metabolism in aortae. Permeabilized vessels were exposed to substrates and inhibitors mimicking lipid metabolism and background oxygen consumption or leak state (state 2), oxidative phosphorylation (+ADP, state 3), maximum oxidative phosphorylation (succinate, state 3S), state 4, and uncoupled respiration (+FCCP) were determined. Respiration rates were normalized to tissue dry weight (n=7-8). Interaction effect ^a^p<0.05, effect of temperature ^b^p<0.05, effect of sex ^c^<0.05, effect of temperature ^†^p=0.08, two-way ANOVA. Data are mean ± SEM.

|  | RT |  | TN |  |
| --- | --- | --- | --- | --- |
| **O<sub>2</sub> pmol/s\*mg** | **Female** | **Male** | **Female** | **Male** |
| **State 3S<sup>b</sup>** | 28.39 ± 2.47 | 25.25 ± 2.12 | 21.03 ± 2.00 | 22.01 ± 0.52 |
| **State 4<sup>b</sup>** | 21.98 ± 2.25 | 20.58 ± 2.12 | 14.58 ± 1.66 | 16.08 ± 1.02 |
| **Uncoupled<sup>b</sup>** | 45.46 ± 3.23 | 38.14 ± 1.64 | 32.60 ± 3.29 | 33.19 ± 0.91 |
| **RCR<sup>b</sup>** | 1.31 ± 0.04 | 1.24 ± 0.03 | 1.46 ± 0.05 | 1.39 ± 0.07 |

### Male aorta had significantly higher expression of antioxidant enzyme MnSOD

In PVAT, MnSOD expression was significantly lower in TN-housed animals as compared with RT-housed animals (p<0.05, Figure 6A). In male aortae, MnSOD expression was significantly elevated in comparison to female rat aortae (p<0.05, Figure 6B).

**Figure 6A and B.**
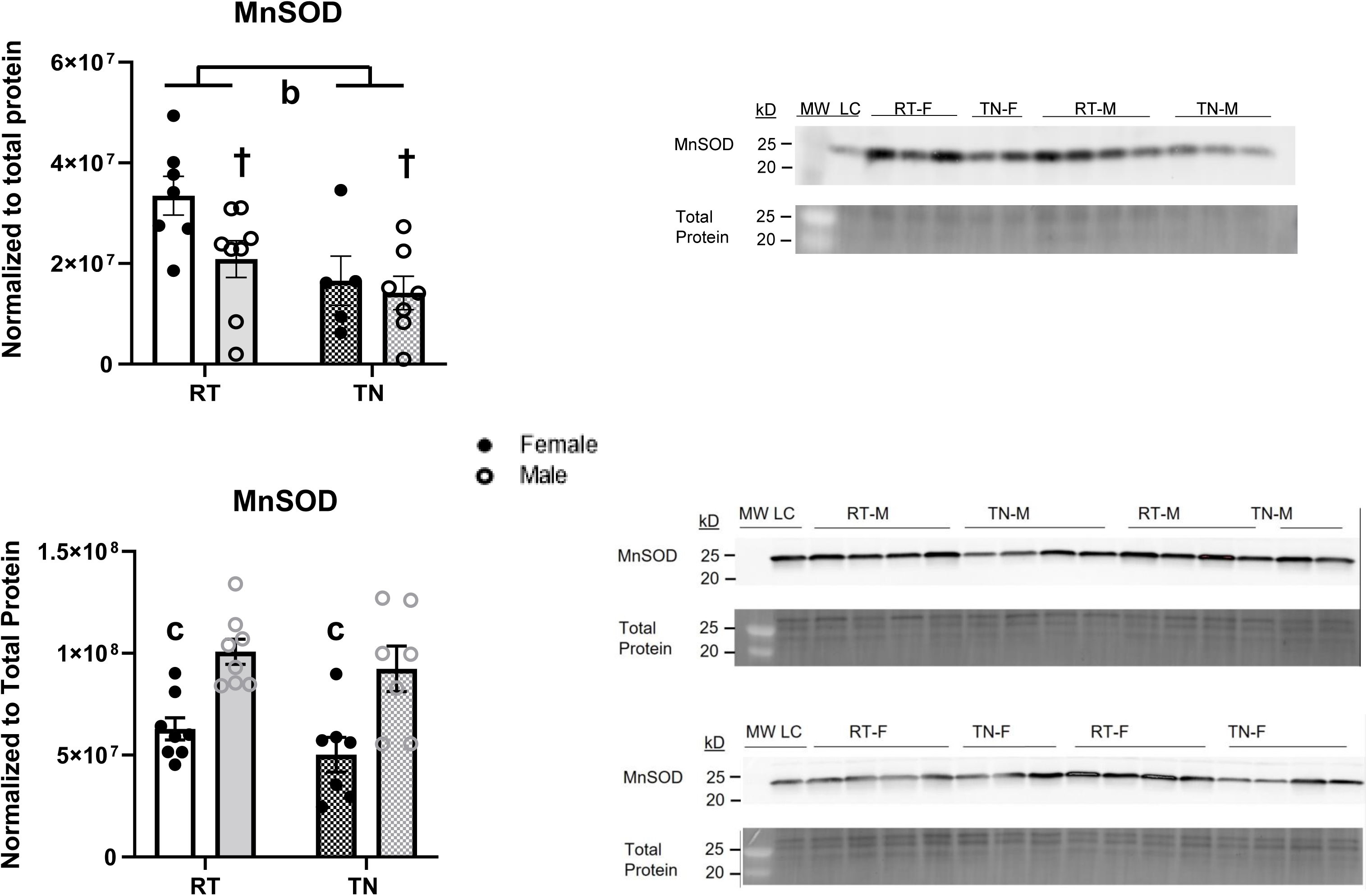
PVAT (A) and aortae (B) protein expression of endogenous antioxidant defenses MnSOD. PVAT and aortae tissue was processed for protein analysis via Western blot analysis for MnSOD (n=7-8). Interaction effect ^a^p<0.05, effect of temperature ^b^p<0.05, effect of sex ^c^<0.05, ^†^p=0.08, two-way ANOVA. Data are mean ± SEM.

## Discussion

We previously reported diminished vasodilation and enhanced vasoconstriction in rats housed at thermoneutrality (27). Here, we further these findings and support our hypothesis that phenotypic changes in PVAT contribute to alterations in vasoreactivity in conduit vessels. We show that TN housing causes PVAT whitening, in turn impacting PVAT’s regulation of vasoreactivity. Vessel response is altered by TN housing, with increased deleterious effects in females. In female aortae, the diminished vasoreactivity seems initially driven by aortae response to temperature; this dysfunction can be restored by exposure to PVAT from room temperature aortae, intriguingly not observed in males. Aortae from animals housed at TN displayed the highest amount of stiffness, prominent in females, consistent with the sex-differences in vasoreactivity. We demonstrate altered signaling upstream of mitochondrial respiration and vasoreactivity in both PVAT and aortae. Taken together, these data support a model of PVAT whitening, leading to diminished PVAT/vascular crosstalk and aorta mitochondrial function.

A main finding of our study shows that PVAT has a sex-specific regulatory role on vascular function, with manifestations at the functional structural and cellular levels. We show that the TN female aortae vasodilation response to SNAP is less than observed in males, suggesting TN-driven sex differences in aortic smooth muscle response; however, we noted significant stiffening in female aorta from TN-housed animals, suggesting that the dilation response in smooth muscle remains intact. These data align with and add to previous reports (10–15). For example, in LDL and *Apoe* knockout mouse models, atherosclerosis progresses more severely in females than males (35, 36). In humans, brachial artery flow-mediate dilation was found to worsen across the menopause transition, illustrating declining endothelial function correlating with diminishing estrogen concentrations (7). Estrogen is known to support eNOS function and NO signaling to vasodilation (4, 6). As peNOS and eNOS expression in our study show only minor sex differences, we hypothesize that estrogen’s impact on eNOS/NO is blunted by TN-housing, currently explored in our laboratory.

eNOS, active in the endothelium, is a main instigator of vasodilation and is compromised in the context of CVD (37, 38). (39); our findings of diminished eNOS expression in aortae from rats housed at TN support a model whereby PVAT whitening and resultant loss of adiponectin, itself a vasodilator and known to be upstream of eNOS (40), could contribute to the diminished eNOS activity in the vasculature. We noted no impact of TN PVAT on RT aortae, potentially due to low concentrations of adiponectin and eNOS activity. We restored vasodilation in TN aorta with RT PVAT, expected if physiological adiponectin signaling to eNOS is present. These data align with other studies showing eNOS and adiponectin as main mechanisms of PVAT/vascular crosstalk (16, 17, 41).

We observe temperature–induced dampening in our mitochondrial respiration measurements in aortae that align with diminished protein expression of complexes I, II and IV. We also observe sex differences in mitochondrial protein expression and MnSOD at the cellular level, consistent with differential regulation of function at the tissue level. As no temperature-related changes in PGC1-α expression were noted, these effects are likely independent of mitochondrial biogenesis. In *ex vivo* mitochondrial respiration experiments, we may be observing a transformation in fatty acid metabolism and/or mitochondrial adaptation to available substrates in aortae from TN animals, as reported in other contexts (11, 13, 42). Additionally, decreased mitochondrial activity at TN may indicate alterations of cellular function subsequent to increased lipid size and storage (43). BAT shows an increase in ATP production when mitochondria are in contact with lipids (43); if BAT is decreasing at TN, this might explain our observations of diminished mitochondrial activity in the presence of lipid and other substrates.

Intriguingly, both male and female animals housed at TN weighed significantly less than those at RT. This is in contrast to other studies observing increased body weight gain in rodents exposed to cold temperatures from either RT housing or TN housing (44, 45). Some studies show no difference in body weight between rats housed at RT and TN (46), while others show increased diet-dependent adiposity at TN temperatures (47). In our previous study utilizing housing temperature to induce vascular dysfunction, we report that diet had no impact on vascular function (28). Weight changes related to TN housing appear to be more complex than just adipose whitening, thus further investigation of potential mechanisms is necessary.

Our study has several limitations. First, we are analyzing PVAT at a specific location, surrounding the thoracic aorta, and our samples keep the stromal vascular fraction intact. This restricts our ability to the PVAT/adventitial cell of origin for the peNOS. The PVAT aortic may behave differently than the mesenteric PVAT depot, for instance. Ongoing studies examine the carotid and mesenteric vascular beds. Secondly, we show major differences in ex vivo aorta response to ACh in intact aorta compared to aorta cleaned of PVAT. We interpret this as the visualization of PVAT regulation of the vessel as compared with the vessels’ maximal vasodilation. This observation agrees with multiple prior reports with PVAT removed (52–54). We acknowledge that we are testing these phenomena ex situ, and the actual changes in aorta diameter, as controlled by PVAT, are a model and may differ from in vivo regulation. Despite these shortcomings, we show a convincing and repeatable model for testing vascular dysfunction with a simple TN housing intervention. We show robust vascular dysfunction that is repaired by PVAT transplant. Our model is ideal for specific vascular questions and sets the stage for hemodynamic cardiac investigations.

Our study is relevant to human physiology in the following ways. First, it has been strongly suggested that human PVAT also contains a mix of BAT and WAT phenotype, including PVAT paracrine signaling to the vasculature (55). Reports indicate that human BAT depots respond to cold exposure by elevating glucose uptake and whole-body energy expenditure (56). In summary, although studies on temperature impacts on human PVAT are uncommon, it follows that BAT in PVAT would respond to either cold or hot temperature exposure. Second, a recent study demonstrates differences in BAT activation between those living in tropical versus temperate locations (57). This implies that human PVAT may have differing paracrine activity depending on climate. Our study supports future work investigating the impact on environmental temperature on human PVAT and vascular crosstalk.

In conclusion, we show that TN mediated impairments in vasoreactivity occur in the context of PVAT whitening in a sex specific manner. This results in a failure of physiological crosstalk between the adipose and vascular tissue, as seen by 1) the RT PVAT exposure to TN aortae normalizing vasoreactivity, only in female aortae, 2) the whitening of TN PVAT resulting in diminished vasodilator adiponectin concentrations and eNOS activity in TN female aortae, and 3) arterial stiffness is higher in TN female aortae concurrent with lower mitochondrial respiration in both male and female TN aortae. Our study contributes valuable insight into the progression of CVD pathology by characterizing PVAT whitening and subsequent regulation of vascular reactivity.

## Acknowledgements

The authors wish to thank Ms. Teri Armstrong and Ms. Melissa Blatzer for their kind assistance with the in vivo measurements. We also thank Mr. Jeremy Rahkola for technical prowess.

## Funding sources

The authors wish to acknowledge the following funding sources: NIH/NCRR CCTSI UL1 RR025780, VA Merit (JEBR BX002046), R01 DK124344-01A1 (JEBR), VA CDA2 (ACK BX003185), Denver Research Institute, and the Ludeman Family Center for Women’s Health Research at the University of Colorado Anschutz Medical Campus Junior Faculty Research Development Grant (ACK).

## Author contributions

ACK and JEBR conceived and designed research, ACK, MMH, JHC, LAK, and GBP performed experiments, ACH, MMH, JHC, GEJ, analyzed data, ACK, KSH, RS, LAW, and JEBR interpreted results of experiments, ACK, JHC, and MMH prepared figures, MMH and ACK drafted manuscript, ACK, MMH, and JEBR edited and revised manuscript, and ACK and JEBR approved final version of the manuscript.

## Data availability

Any data is freely available upon request to the corresponding author.

## Conflict of Interest/Disclosures

There are no conflicts of interest with any of the authors, and no relevant disclosures.

## New and Noteworthy

Perivascular adipose tissue (PVAT) regulates vascular reactivity, and this crosstalk involves adipokines and cellular signaling. PVAT regulation may be altered in cardiovascular disease. We report that housing animals at thermoneutrality alters PVAT phenotype from BAT to WAT. These temperature-induced alterations are associated with compromised vasoreactivity with sex-specific changes in PVAT and aortae. As CVD pathology differs between the sexes, our results illustrate sex-dependent differences in vascular adaptation to stress.

## Supplemental Figure Legends

**S1A: Representative blots of Figures 3, 4A and 4B.**

**S1B: Whole blots depicting representative images used for Figures 3, 4A and 4B.**

**S2A: Representative blots of Figures 4C and 4D**.

**S2B: Whole blots depicting representative images used for Figures 4C and 4D**.

**S2C: Representative blots of Figures 6A**.

**S2D: Whole blots depicting representative images used for Figures 6A**. Blots for PGC-1α have been cut originally at 75 kD.

**S2E: Representative blots of Figures 6B**.

**S2F: Whole blots depicting representative images used for Figures 6B**.

**S3A: Representative blots of Figures 7A and B**.

**S3B: Whole blots depicting representative images used for Figures 7A and 7B**. Blots for SOD and MnSOD have been originally at 50 kD.

**Supplemental Table 1: Mitochondrial respiration, carbohydrate SUIT protocol.** Permeabilized vessels were exposed to substrates and inhibitors mimicking lipid metabolism and background oxygen consumption or leak state (state 2), oxidative phosphorylation (+ADP, state 3), maximum oxidative phosphorylation (succinate, state 3S), state 4, and uncoupled respiration (+FCCP) were determined. Respiration rates were normalized to tissue dry weight (n=7-8). Interaction effect ^a^p<0.05, effect of temperature ^b^p<0.05, effect of sex ^c^<0.05, effect of temperature ^†^p=0.08, two-way ANOVA. Data are mean ± SEM.

**Supplemental Table 2: List of antibodies’ suppliers, catalog numbers, lot numbers and concentrations.**

## Notes

### Competing Interest Statement

The authors have declared no competing interest.

