## Supplemental Figures for "Perivascular adipose tissue remodeling impairs vasoreactivity in thermoneutral-housed rats"

### Slide 1
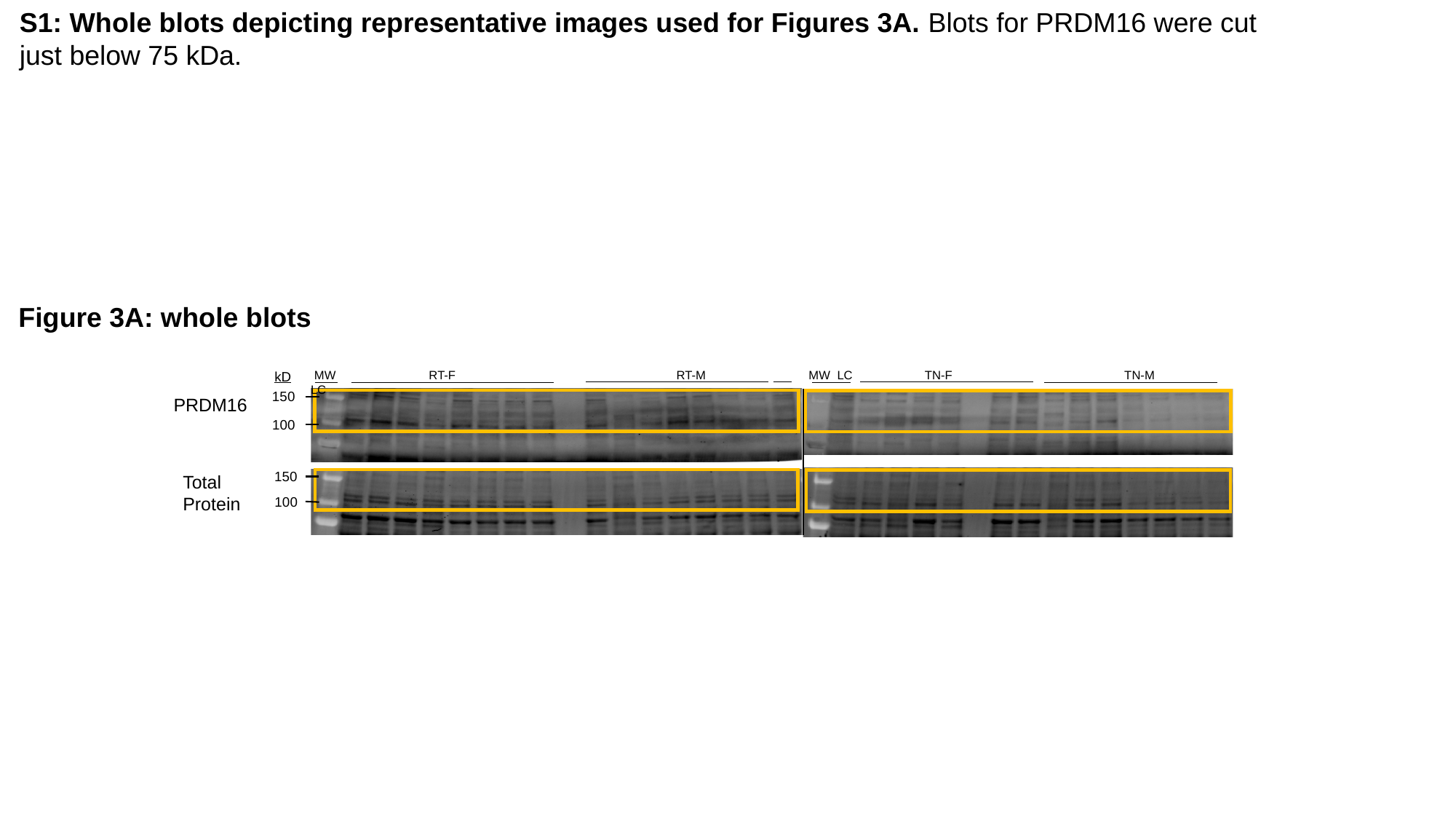

S1: Whole blots depicting representative images used for Figures 3A. Blots for PRDM16 were cut just below 75 kDa.
Figure 3A: whole blots
MW LC TN-F TN-M
 MW RT-F RT-M LC
kD
150
PRDM16
100
150
Total Protein
100

### Slide 2
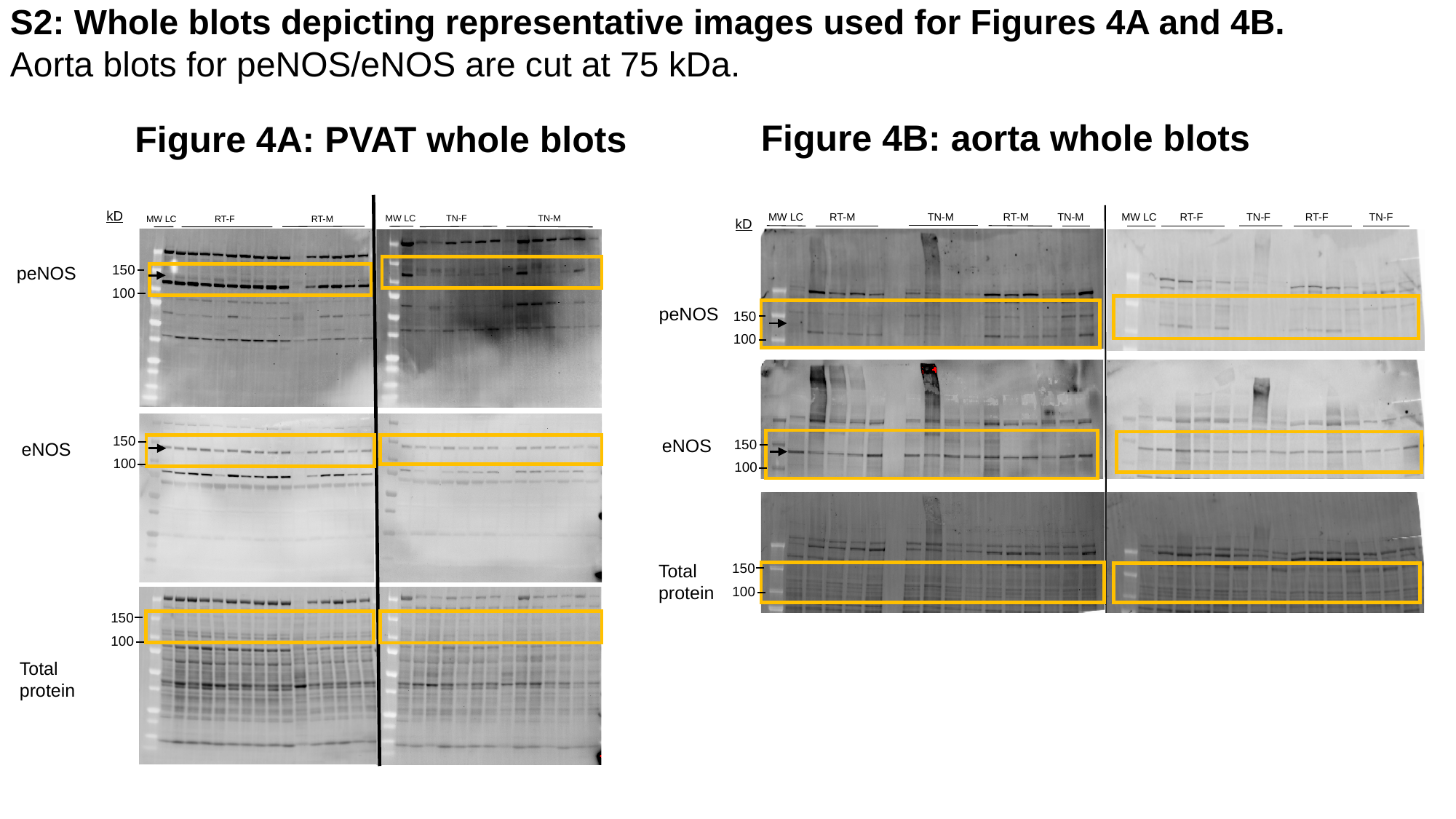

S2: Whole blots depicting representative images used for Figures 4A and 4B. Aorta blots for peNOS/eNOS are cut at 75 kDa.
Figure 4B: aorta whole blots
Figure 4A: PVAT whole blots
kD
MW LC            TN-F                            TN-M
MW LC               RT-F                              RT-M
150
peNOS
100
150
eNOS
100
150
100
MW LC        RT-F               TN-F            RT-F              TN-F
MW LC RT-M TN-M RT-M TN-M
kD
peNOS
150
100
eNOS
150
100
150
Total protein
100
Total protein

### Slide 3
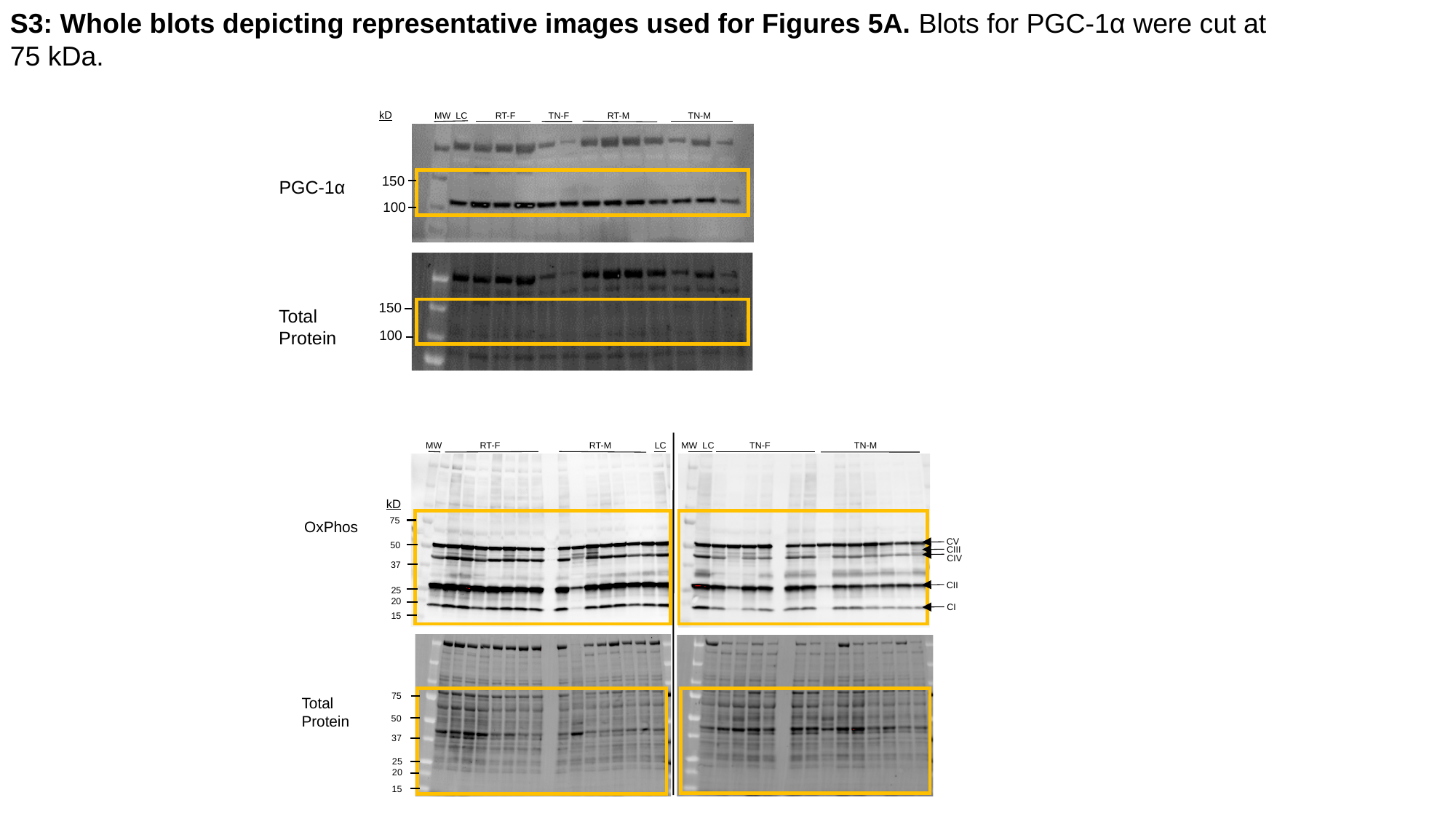

S3: Whole blots depicting representative images used for Figures 5A. Blots for PGC-1α were cut at 75 kDa.
kD
MW  LC           RT-F             TN-F               RT-M                       TN-M
150
PGC-1α
100
150
Total Protein
100
MW  LC              TN-F                                 TN-M
 MW               RT-F                                   RT-M                 LC
kD
75
OxPhos
CV
50
CIII
CIV
37
CII
25
20
CI
15
75
Total
Protein
50
37
25
20
15

### Slide 4
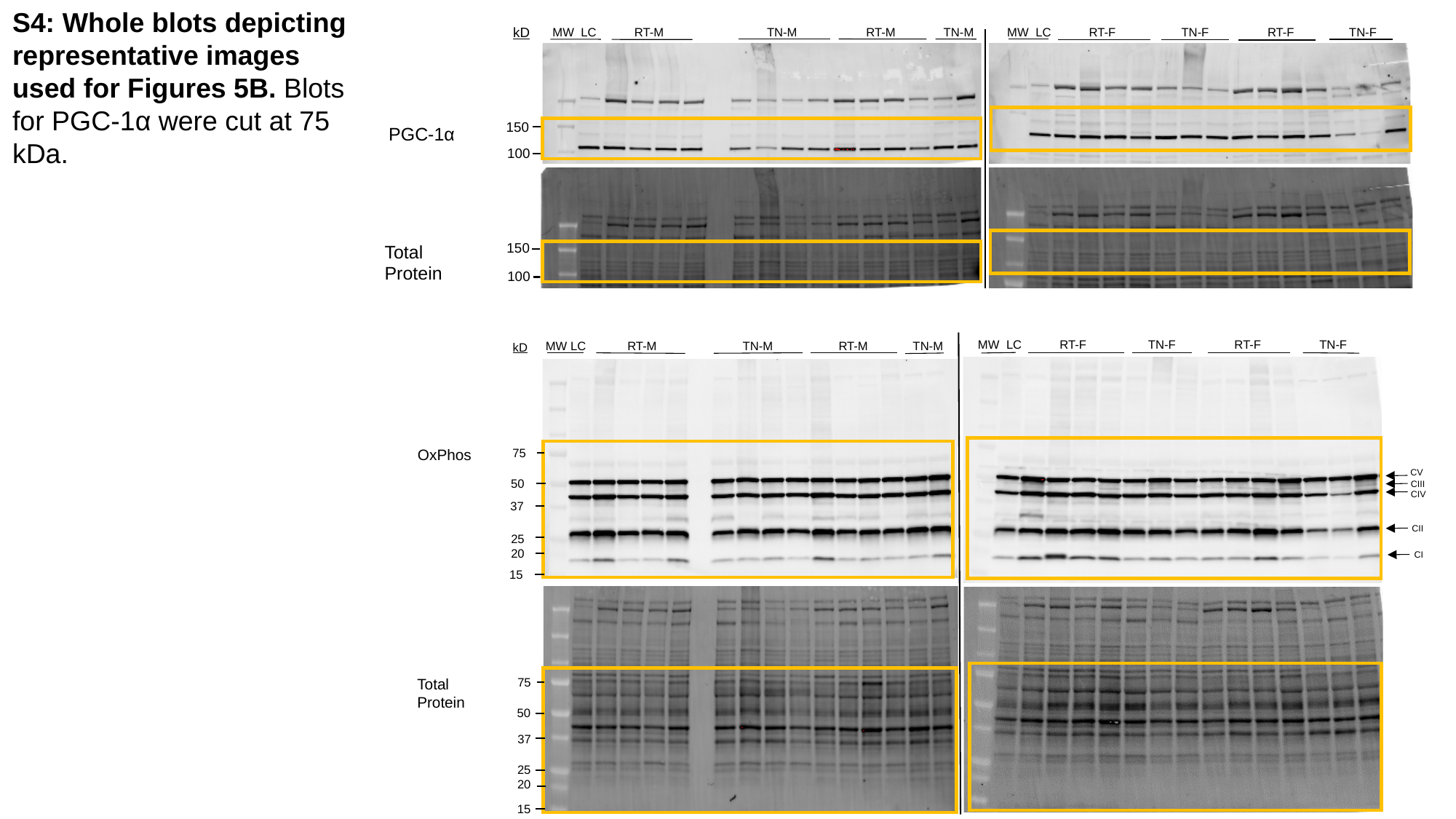

S4: Whole blots depicting representative images used for Figures 5B. Blots for PGC-1α were cut at 75 kDa.
kD
MW  LC           RT-M                              TN-M                    RT-M              TN-M
MW  LC           RT-F                   TN-F                 RT-F                TN-F
150
PGC-1α
100
150
Total Protein
100
MW  LC           RT-F                  TN-F                 RT-F                 TN-F
 MW LC            RT-M                         TN-M                   RT-M             TN-M
kD
OxPhos
75
50
37
25
20
15
Total
Protein
75
50
37
25
20
15
CV
CIII
CIV
CII
CI

### Slide 5
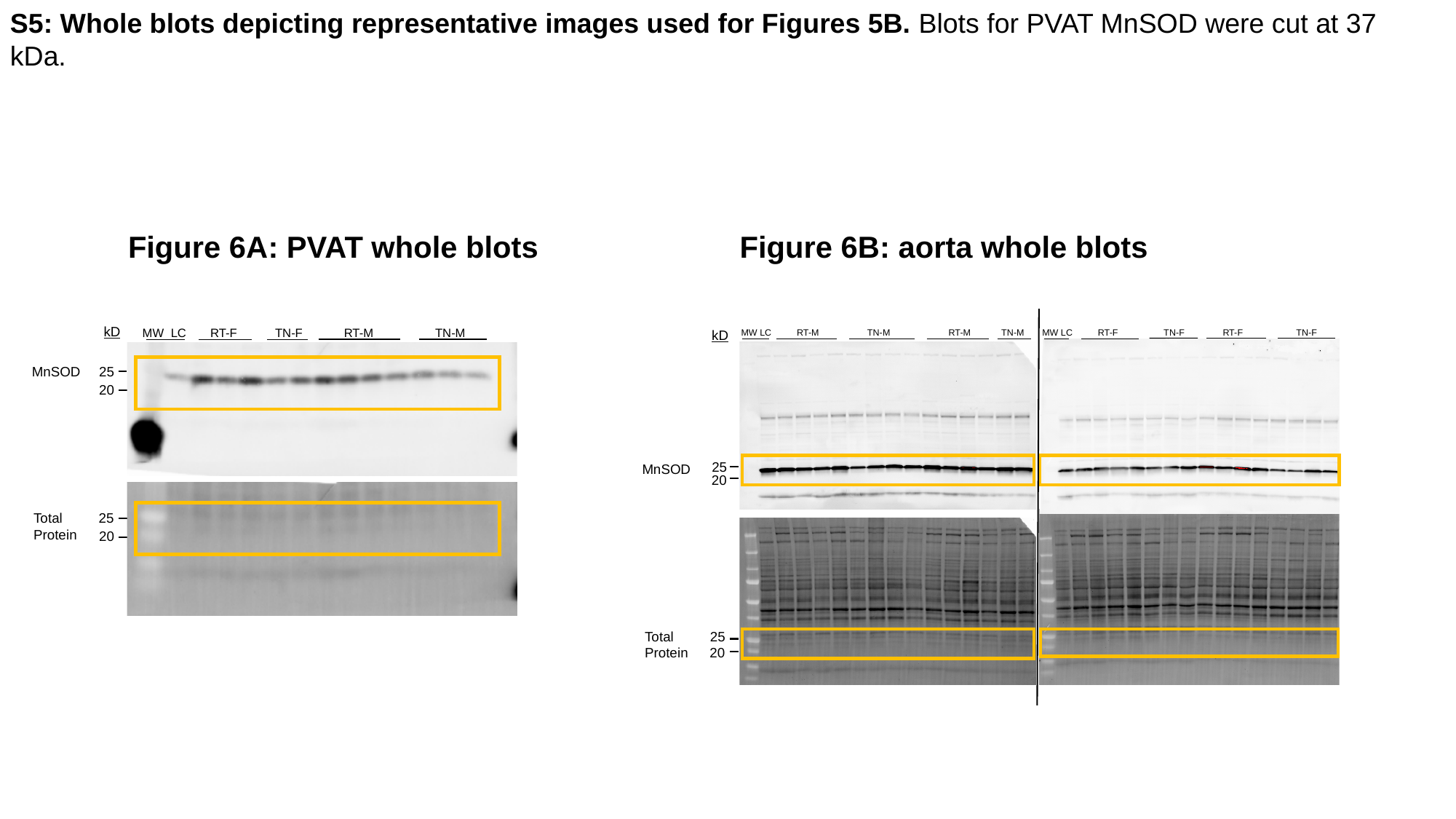

S5: Whole blots depicting representative images used for Figures 5B. Blots for PVAT MnSOD were cut at 37 kDa.
Figure 6B: aorta whole blots
Figure 6A: PVAT whole blots
MW LC          RT-F                  TN-F               RT-F                     TN-F
kD
MW LC          RT-M                   TN-M                       RT-M            TN-M
25
MnSOD
20
Total
Protein
25
20
kD
MW  LC       RT-F           TN-F            RT-M                  TN-M
25
MnSOD
20
25
Total
Protein
20
