## Supplemental Table 1 for "Perivascular adipose tissue remodeling impairs vasoreactivity in thermoneutral-housed rats"

**Supplemental Table 1:** Permeabilized vessels were exposed to substrates and inhibitors mimicking carbohydrate metabolism and background oxygen consumption or leak state (state 2), oxidative phosphorylation (+ADP, state 3), maximum oxidative phosphorylation (succinate, state 3S), state 4, and uncoupled respiration (+FCCP) were determined. Respiration rates were normalized to tissue dry weight (n=7-8). Interaction effect ^a^p<0.05, effect of temperature ^b^p<0.05, effect of sex ^c^<0.05, effect of temperature ^†^p=0.08, two-way ANOVA. Data are mean ± SEM.

|  | **RT** | | **TN** | |
| --- | --- | --- | --- | --- |
| **O_2_ pmol/s*mg** | **Female** | **Male** | **Female** | **Male** |
| **State 3S†** | 35.95 ± 6.20 | 28.24 ± 3.19 | 26.88 ± 2.06 | 21.25 ± 1.40 |
| **State 4^b^** | 21.98 ± 2.25 | 21.69 ± 2.68 | 14.58 ± 1.66 | 15.11 ± 1.35 |
| **Uncoupled** | 45.46 ± 3.27 | 60.15 ± 7.14 | 32.60 ± 3.29 | 50.15 ± 2.43 |
| **RCR** | 1.34 ± 0.05 | 1.32 ± 0.06 | 1.41 ± 0.04 | 1.43 ± 0.05 |
