## Supplemental Table 2 for "Perivascular adipose tissue remodeling impairs vasoreactivity in thermoneutral-housed rats"

**Supplemental Table 2: List of antibodies’ suppliers, catalog numbers, lot numbers and concentrations.**

|  | **Type** | **Name** | **Source** | **Dilution** | **Secondary Antibody** | **Manufacturer** | **Catalog** | **Lot:** | **RRID** | **Application** |
| --- | --- | --- | --- | --- | --- | --- | --- | --- | --- | --- |
| **1** | Primary | phospho-AMPKa (Thr172) Antibody | Rabbit | 1:500 | 12 | Cell Signaling Technology | 2531S | 16 | AB_330330 | Western Blot |
| **2** | Primary | AMPK (F6) Mouse mAb | Mouse | 1:500 | 13 | Cell Signaling Technology | 2793S | 8 | AB_915794 | Western Blot |
| **3** | Primary | phospho-eNOS (Ser1177) Polyclonal | Rabbit | 1:500 | 12 | Thermo Fisher Scientific | PA5-17917 | YD3887 188A | AB_10986074 | Western Blot |
| **4** | Primary | eNOS (6H2) Mouse mAb | Mouse | 1:500 | 13 | Cell Signaling Technology | 5880S | 2 | AB_10850618 | Western Blot |
| **5** | Primary | SOD2 (D9V9C) | Rabbit | 1:500 | 14 | Cell Signaling Technology | 13194S | 1 | AB_2750869 | Western Blot |
| **6** | Primary | Anti-Superoxide Dismutase 1 Antibody | Rabbit | 1:1000 | 14 | abcam | ab13498 | GR80256-81 | AB_300402 | Western Blot |
| **7** | Primary | PGC1a | Rabbit | 1:1000 | 14 | abcam | ab54481 | GR3384581-1 | AB_881987 | Western Blot |
| **9** | Primary | Total OXPHOS Rodent WB Antibody Cocktail | Mouse | 1:1000 | 13 | abcam | ab110413 | 2101009677 | AB_2629281 | Western Blot |
| **10** | Primary | PRDM16 Polyclonal Antibody | Rabbit | 1:500 | 12 | Invitrogen | 720206 | 2448891 | AB_2664823 | Western Blot |
| **11** | Primary | UCP1 Polyclonal Antibody | Rabbit | 1:125 | 16 | Thermo Fisher Scientific | pa5-29575 | we3290611B | AB_2547051 | IHC |
| **12** | Secondary | IRDye 800CW goat anti-rabbit | Goat | 1:10,000 | -------- | Li-Cor | 926-32211 | unknown | AB_621843 | Western Blot |
| **13** | Secondary | Goat Anti-Mouse IgG Starbright Blue 700 | Goat | 1:5,000 | -------- | Bio-Rad Laboratories | 12004159 | 64433886 | AB_2884948 | Western Blot |
| **14** | Secondary | Goat Anti-Rabbit IgG Starbright Blue 700 | Goat | 1:5,000 | -------- | Bio-Rad Laboratories | 12004161 | 64484700 | AB_2721073 | Western Blot |
| **15** | Congugated | wheat germ agglutinin, alexa fluor 555 conjugate | --------- | 1:5 | -------- | Thermo Fisher Scientific | W32464 | 2110867 | ----------- | IHC |
| **16** | Secondary | Goat Anti-Rabbit IgG (H&L) Cross-Absorbed Secondary Antibody, Cyanine5 | Goat | 1:500 | -------- | Invitrogen | a10523 | 2415914 | AB_2534032 | IHC |
